# Semantic-Aware Graph Embedding Approach Uncovers LC-61, a Potent Anti–*Leishmania infantum* Compound

**DOI:** 10.1101/2025.11.19.687645

**Authors:** Vinícius Alexandre Fiaia Costa, Alexandra Maria dos Santos Carvalho, Rafael Consolin Chelucci, Felipe da Silva Mendonça de Melo, Gustavo Santos Sandes Felizardo, Clarissa Alves Carneiro Bernardes, Holli-Joi Martin, Rodolpho de Campos Braga, Sébastien Charneau, Eugene N. Muratov, Adriano Defini Andricopulo, Izabela Marques Dourado Bastos, Bruno Junior Neves

**Affiliations:** Laboratory of Cheminformatics, Faculty of Pharmacy, Federal University of Goiás, Goiânia, Brazil; Pathogen-Host Interface Laboratory, Department of Cell Biology, Institute of Biological Sciences, University of Brasilia, Brasilia, Brazil; Laboratory of Medicinal and Computational Chemistry, Sao Carlos Institute of Physics, University of Sao Paulo, IFSC – USP, 13566-590, Sao Carlos, SP, Brazil; Laboratory for Molecular Modeling, UNC Eshelman School of Pharmacy, University of North Carolina at Chapel Hill, North Carolina, USA; Laboratory of Protein Chemistry and Biochemistry, Department of Cell Biology, Institute of Biological Sciences, University of Brasilia, Brasilia, Brazil; InsilicAll Ltda, São Paulo, Brazil

**Keywords:** Visceral leishmaniasis, drug discovery, graph neural networks, embeddings, and counterfactuality

## Abstract

Visceral leishmaniasis caused by *Leishmania infantum* remains a lethal disease with few therapeutic options, necessitating innovative computational methods approaches to accelerate drug discovery. Here, we present a semantic-aware graph neural network (GNN) framework that features holistic mechanisms to capture long-range molecular interactions and the chemical semantics of antileishmanial compounds. Across two classificatory antileishmanial datasets, our holistic GNNs demonstrated significant improvements in predictive performance, with area under the receiver operating characteristic curve (AUROC) increases of 2.2–29.2% on the unbalanced dataset (1 µM threshold) and 3.4–22.5% on the balanced dataset (10 µM threshold) compared to default GNNs. Subsequently, the framework was applied to screen a library of approximately 1.3 million compounds, pinpointing LC-61 as a potent antileishmanial agent with nanomolar activity against intracellular *L. infantum* (IC_50_ = 0.076 µM) and minimal cytotoxicity to macrophages (THP-1 CC_50_ = 157 µM). A comprehensive *in vitro* ADME profiling revealed that LC-61 combines high solubility at both acidic and physiological pH (>28 µg/mL), balanced lipophilicity (eLogD = 4.07), and favorable passive permeability (PAMPA = 4.86 × 10^-6^ cm/s), while exhibiting lower microsomal stability. Overall, our semantic-aware GNN framework effectively accelerated the discovery of LC-61, a novel and biologically validated hit suitable for hit-to-lead optimization.

## 1. INTRODUCTION

Visceral leishmaniasis (VL) is a potentially fatal vector-borne disease transmitted by the bite of infected sandflies. It is primarily caused by two protozoan parasites of the *Leishmania* genus: *Leishmania donovani*, which is predominantly found in Africa and Asia, and *Leishmania infantum*, which is mainly present in Latin America and the Middle East.^1–3^ This disease affects vital organs such as the liver, spleen, and bone marrow, manifesting clinically as weight loss, fatigue, splenomegaly, hepatomegaly, anemia, and irregular fever.^4^ According to the World Health Organization, most cases of VL are concentrated in Brazil, East Africa, and India. Although it is estimated that 50,000–90,000 new cases occur globally each year, only 25–45% of these are officially reported, highlighting significant gaps in disease surveillance.^5^

In the absence of an effective vaccine for humans, disease treatment primarily relies on a limited pharmacological arsenal, including amphotericin B, miltefosine, and pentavalent antimonials. However, these therapeutic options come with significant drawbacks, such as high costs, complex long-term treatment protocols, and severe adverse effects, including nephrotoxicity, cardiotoxicity, and hepatotoxicity. The clinical utility of these agents is further constrained by emerging drug resistance and the requirement for parenteral administration in resource-limited settings. Consequently, there is a pressing need for innovative drug discovery approaches to expand the therapeutic landscape for this neglected tropical disease.^6–8^

Identifying antileishmanial hits poses a significant challenge in early-stage drug discovery, particularly due to the time and resources required for experimental screening for efficacy and safety. Over recent decades, artificial intelligence (AI)-driven methodologies have revolutionized this process by significantly enhancing both the efficiency and cost-effectiveness of molecular property predictions.^9,10^ One of the core challenges in AI-based drug discovery is accurately representing chemical structures. Earlier machine learning approaches relied on simplistic representations using handcrafted features like molecular descriptors and fingerprints.^11–13^ Molecular descriptors quantify the physical and chemical attributes of compounds,^14^ while fingerprints encode the presence or absence of specific molecular substructures as bit vectors.^15^ Although these traditional methods have contributed significantly to previous efforts, they are inherently limited in their capacity to capture the nuanced structural and topological complexity of molecules, thus highlighting the need for more advanced and expressive representations in contemporary AI-driven drug discovery.^16,17^

Graph Neural Networks (GNNs) significantly advance this area, offering highly adaptable architectures and feature representations for modeling complex biological properties, such as antileishmanial activity.^18,19^ Unlike traditional deep learning models that represent molecules as linear sequences (e.g., SMILES strings) or bit vectors (e.g., molecular fingerprints), GNNs operate directly on molecular graphs, where atoms are represented as nodes and chemical bonds as edges. Formally, a molecular graph is defined as 𝐺 = (𝑉, 𝐸), with 𝑉 denoting the set of atoms and 𝐸 the set of bonds. Node features, such as atom types and hybridization states, are encoded in a feature matrix 𝑋, where each node 𝑣 ∈ 𝑉 is associated with an initial vector 𝑥_𝑣_ ∈ ℝ^𝑑^. Bond attributes are similarly represented in an edge feature matrix. The connectivity of the molecule is captured by an adjacency matrix 𝐴, where 𝐴_𝑖𝑗_ = 1 indicates a bond between atoms 𝑣_𝑖_ and 𝑣_𝑗_.^20^

A wide range of GNN architectures—such as Message Passing Neural Networks (MPNN),^21^ Graph Attention Networks (GAT),^22^ Graph Isomorphism Networks (GIN),^23^ and Attentive Fingerprint (AttentiveFP)^24^—have been proposed and continually refined through the integration of sophisticated mechanisms. These include attention schemes,^25,26^ structure-aware update rules,^27,28^ hierarchical representation learning,^29,30^ and high-order relational modeling.^31^ In general, GNNs operate through a message-passing paradigm, wherein node features are iteratively updated by aggregating information from their immediate neighbors and, optionally, from edge features, using learnable functions that capture local chemical environments.^19^ This process facilitates the incorporation of both local and, to some extent, global structural context, ultimately enhancing the encoding of structural and contextual information and resulting in improved performance across diverse cheminformatics tasks.

Despite substantial progress in molecular representation learning, GNNs encounter fundamental limitations when applied to complex drug discovery tasks. Conventional message-passing architectures operate predominantly on local neighborhoods, which restricts the modeling of long-range dependencies essential for capturing global pharmacophoric interactions.^32,33^ As network depth increases, these models are also prone to over-smoothing, where node embeddings become indistinguishable, and over-squashing, where information from distant nodes is inadequately compressed, thereby impairing the expressivity and semantic richness of the learned representations.^34–36^ Consequently, there is a pressing need for holistic GNN frameworks that can effectively model extended molecular interactions while preserving chemical semantics relevant to antiparasitic activity.

This study addresses these fundamental limitations through the development of a holistic, semantic-aware GNN-based framework that enhances long-range information propagation and preserves chemical semantics for predicting antileishmanial activity against *L. infantum*. We hypothesized that integrating semantic molecular context into graph learning would improve predictive performance and facilitate the identification of novel antileishmanial compounds. Using curated bioassay data from ChEMBL,^37^ enhanced architecture demonstrated significant improvements in predictive performance compared to standard GNN counterparts in ablation studies. The framework also incorporates a counterfactual explainability module, enabling atom-level interpretation of model outputs and highlighting key substructures responsible for antileishmanial activity. The best-performing GNN models were subsequently employed in a large-scale virtual screening (VS) of 1.3 million commercially available compounds from the ChemBridge database, yielding 18 candidates for biological evaluation. This campaign achieved a 50% experimental hit rate and led to the identification of LC-61, a chemically novel scaffold exhibiting nanomolar antileishmanial potency, low cytotoxicity, and a favorable *in vitro* ADME profile, establishing it as a promising hit-to-lead candidate for further optimization.

## 2. RESULTS & DISCUSSION

### 2.1. Data description

A total of 1,415 compounds with reported *in vitro* antileishmanial activity against intracellular *L. infantum* were compiled from ChEMBL to support the development and evaluation of holistic GNN models. These compounds were subsequently classified as active or inactive using two distinct activity thresholds. A stringent 1 µM threshold was applied to capture higher potency profiles, resulting in 213 actives (IC₅₀ ≤ 1 µM) and 1,220 inactives (IC₅₀ > 1 µM), yielding a highly imbalanced dataset with a lower prevalence of actives (Figure 1a). Additionally, a 10 µM threshold, consistent with the hit criteria proposed by Katsuno et al.,^38^ was applied to generate a more balanced dataset comprising 641 actives (IC₅₀ ≤ 10 µM) and 774 inactives (IC₅₀ > 10 µM) (Figure 1b).

**Figure 1.**
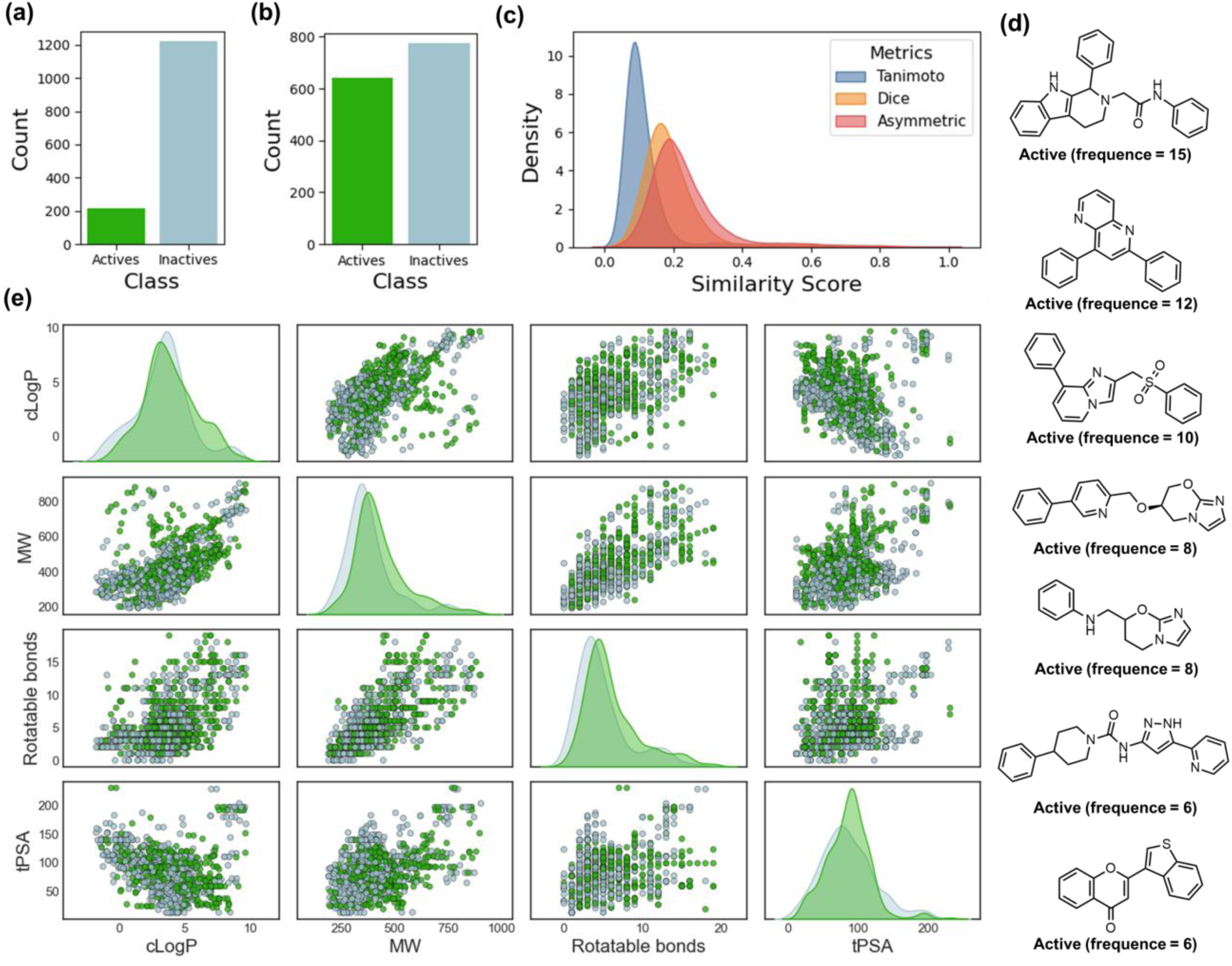
Chemical space analysis and structural profiling of active and inactive compounds used for model training. Distribution of actives and inactives in (**a**) unbalanced and (**b**) balanced datasets developed using 1 µM and 10 µM thresholds, respectively. (**c**) Density plots of pairwise similarity scores between compounds calculated using Tanimoto, Dice, and Asymmetric metrics. (**d**) Molecular structures of representative scaffolds observed among actives, annotated with their frequency of occurrence in the balanced dataset. (**e**) Pairwise scatter plots and kernel density estimates for cLogP, MW, number of rotatable bonds, and tPSA for actives (green) and inactives (gray) from the balanced dataset.

Subsequently, pairwise molecular similarity distributions were computed using ECFP4 fingerprints with Tanimoto, Dice, and Asymmetric metrics, revealing overall low similarity scores across compound pairs and indicating a chemically diverse dataset (Figure 1c). This diversity is essential for minimizing dataset bias and enhancing scaffold-hopping capabilities in predictive pipelines. Furthermore, analysis of Bemis-Murcko scaffolds identified 659 distinct chemotypes across the dataset. Among these, seven privileged scaffolds frequently observed in actives but rarely present in inactives are represented in Figure 1d. These scaffolds predominantly encompass polyaromatic and heterocyclic frameworks commonly enriched in bioactive libraries, suggesting the presence of privileged structures that may drive antileishmanial activity.

Scatter plot matrices and marginal density plots of key molecular descriptors, including calculated logarithm of the partition coefficient (cLogP), molecular weight (MW), number of rotatable bonds, and topological polar surface area (tPSA), were generated to visualize the physicochemical property distribution of active and inactive compounds. As shown in Figure 1e, substantial overlap was observed between actives and inactives across these physicochemical dimensions, underscoring the limitations of conventional property-based thresholds for distinguishing antileishmanial activity in this dataset. These findings highlight the necessity of advanced representation learning approaches capable of capturing non-linear structure–activity relationships and topological patterns beyond classical descriptor-based separations.

### 2.2. Model structure

To address the limitations of both traditional molecular descriptors and conventional message-passing GNNs, a systematic enhancement of MPNN, GAT, and GIN architectures was implemented through structural mechanisms that enable holistic molecular representations (Figure 2a).

**Figure 2.**
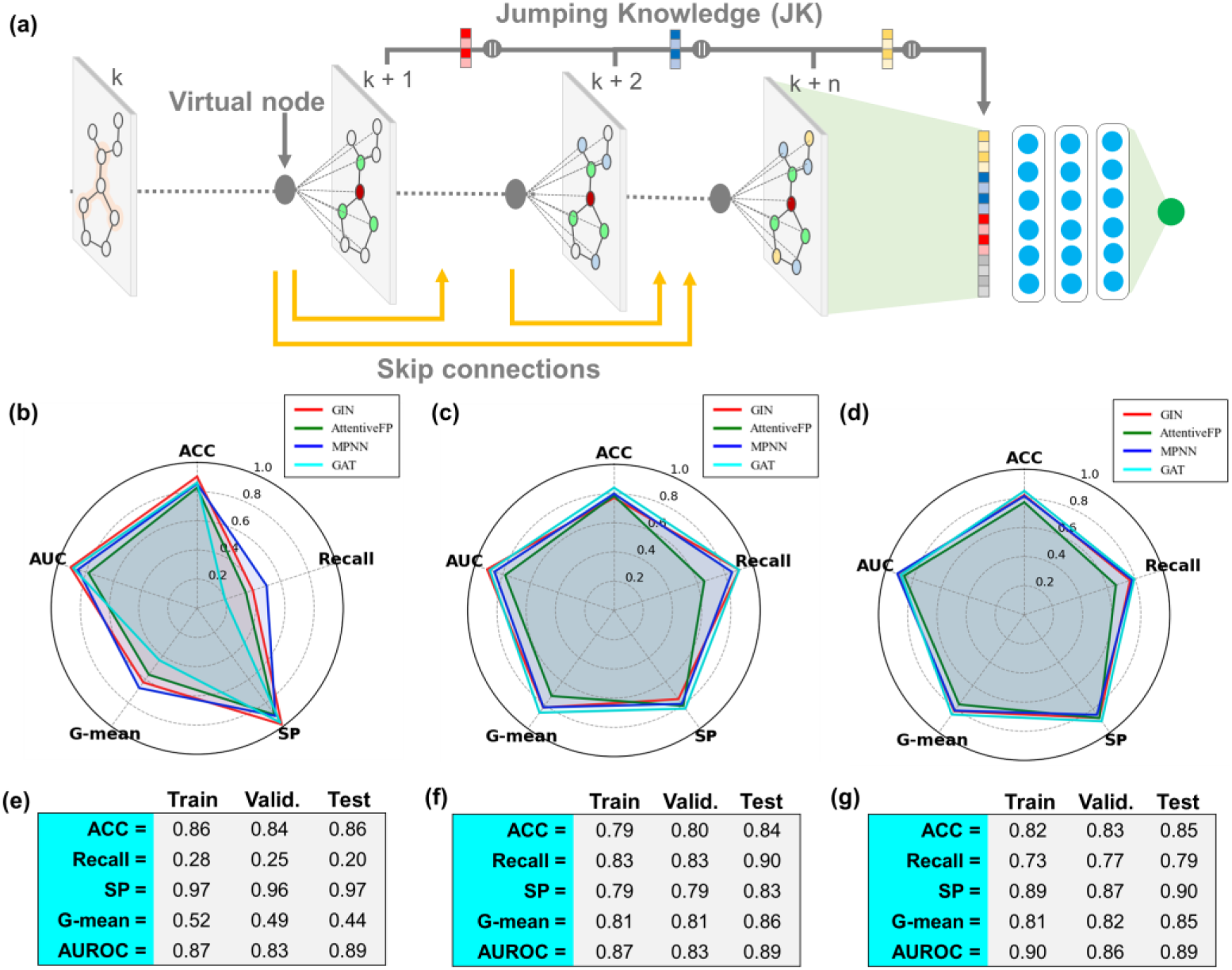
Model architecture and performance evaluation of holistic GNNs for antileishmanial activity prediction. (**a**) Schematic representation of the virtual nodes, skip connections, and the Jumping Knowledge (JK) mechanisms incorporated in GNNs. Radar plots summarizing the performance of holistic GNNs on the unbalanced dataset (**b**) before and (**c**) after calibration based on the empirical distribution of predicted probabilities. (**d**) Radar plots summarize the performance of GNNs for binary classification using the balanced dataset. Detailed statistical results of the GAT developed using the unbalanced dataset (**e**) before and (**f**) after calibration based on the empirical distribution of predicted probabilities. (**g**) Detailed statistical results of the GAT developed using the balanced dataset.

Three key architectural innovations were integrated: First, skip connections were incorporated to mitigate over-smoothing and gradient degradation during deep message-passing, preserving atom-level identity and pharmacophoric signals while ensuring stable training.^39,40^ In parallel, a virtual node was incorporated into each molecular graph to enrich chemical semantics by forging long-range connections among pharmacophoric and structural features, thereby unifying local interactions within a global molecular context.^41^ This strategy strengthens the model’s ability to capture global information; however, it does not fully resolve the challenges arising when relevant signals are distributed across different topological ranges. In view of this, the Jumping Knowledge (JK) mechanism was incorporated into GNN architectures to address this limitation by allowing the model to adaptively aggregate information from multiple message-passing depths, effectively capturing signals at varying neighborhood ranges.^42^

Furthermore, the state-of-the-art AttentiveFP architecture was chosen for this study as baseline because it inherently implements analogous multiscale mechanisms to those introduced: (i) deploying skip connections within each attention layer; (ii) incorporating a global context vector similar to virtual node by broadcasting molecular-level information to every atomic node; and (iii) leveraging a recurrent GRU module that emulates the JK mechanism by adaptively integrating information across past message-passing iterations.

### 2.3. Model performance

The holistic GNN architectures were evaluated using both unbalanced and balanced datasets to assess their predictive capabilities for antileishmanial activity. Initial models developed using the unbalanced dataset demonstrated promising performance, with area under the receiver operating characteristic curve (AUROC) values ranging from 0.78 to 0.891, and G-mean scores ranging from 0.44 to 0.67, reflecting the challenges imposed by class imbalance while confirming the model’s ability to capture patterns associated with antileishmanial activity (Figure 2b and Table S1). To address the classification bias introduced by dataset imbalance, a probability calibration procedure was subsequently performed using the empirical distribution of predicted probabilities, leading to the reclassification of compound predictions based on adjusted decision criteria. This adjustment yielded more balanced recall (0.65–0.90) and specificity (0.75–0.83). Consequently, G-mean scores improved substantially (0.72–0.86), enhancing the practical utility of these models for activity prediction (Figure 2c and Table S2). In contrast, models trained on the balanced dataset exhibited superior baseline performance without calibration, achieving AUROC values of 0.86 –0.91 and G-mean scores of 0.76–0.84 (Figure 2d). This improvement highlights the impact of dataset balance in facilitating the learning of activity patterns and enhancing predictive stability.

Among the evaluated architectures, GAT emerged as the top performer for both thresholds. On the unbalanced dataset, GAT delivered the highest discrimination (Figure 2e-f), reaching a test set accuracy (ACC) = 0.84, recall = 0.90, specificity = 0.83, G-mean = 0.86, and AUROC = 0.89 after probabilistic calibration (probability threshold = 0.13). On the balanced dataset, GAT again ranked first with a test set ACC = 0.85, recall = 0.79, specificity = 0.90, G-mean = 0.84, and AUROC = 0.89 (Figure 2g).

The temporal evolution of latent-space organization during training was visualized with t-SNE projections to track how class separation emerged across epochs (Figure 3). Both models exhibited a progressive increase in separation between active and inactive embeddings, with clusters becoming more compact and inter-class overlap diminishing as training advanced. kernel-density plots reveal clearer bimodality over time, with peak locations drifting apart and overlap diminishing, which is expected under robust representation learning rather than overfitting. These observations confirm the approach’s ability to encode features predictive of antileishmanial activity and demonstrate the stability of the learned representations throughout training.

**Figure 3.**
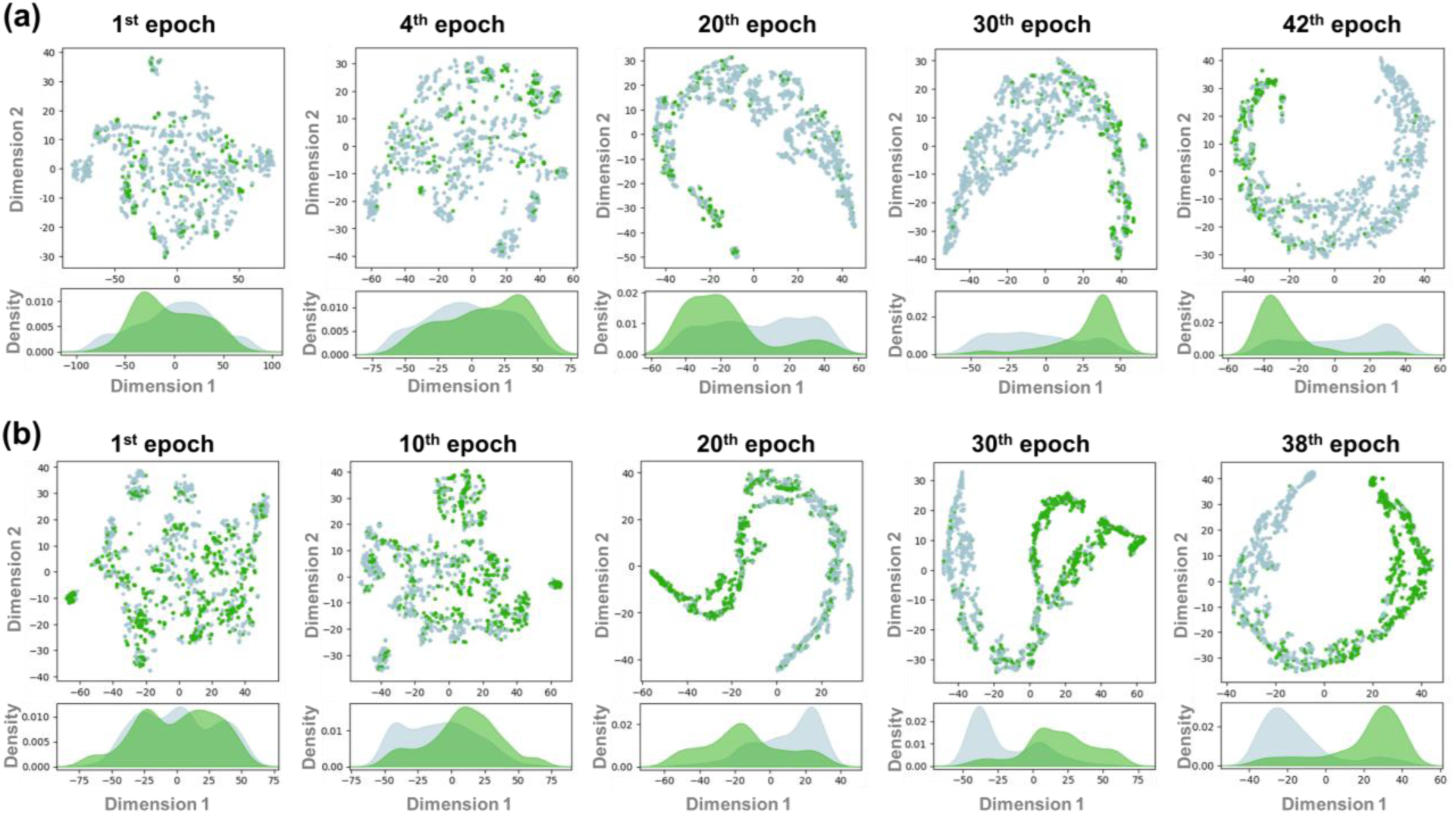
Temporal evolution of latent-space organization during training of GAT. (**a**) t-SNE projections for model trained on the unbalanced dataset; (**b**) t-SNE projections for model trained on the balanced dataset (selected epochs). Green and gray points represent embeddings of active and inactive compounds, respectively. Kernel-density plots of the first t-SNE dimension illustrate the evolving separation between active and inactive embeddings.

### 2.4. Ablation study

A systematic ablation study was conducted to evaluate the individual and combined contributions of holistic components to model performance. Each component (virtual node, skip connections, and JK) was sequentially removed from the holistic GAT architecture, with performance evaluated on both antileishmanial datasets. As shown in Figure 4, the enhanced GAT architecture improved the AUROC by 2.2–29.2% on the unbalanced dataset and by 3.4–22.5% on the balanced dataset. On the unbalanced dataset, removing the virtual node caused a modest decline (–2.2%, 0.89 → 0.87), while omission of skip connections led to a larger reduction (–5.6%, 0.89 → 0.84). The absence of JK had the most pronounced effect, lowering AUROC by 16.9% (0.89 → 0.74). When both skip connections and JK were removed simultaneously, performance decreased by 13.5% (0.89 → 0.77), and the complete removal of all mechanisms yielded the largest drop (–29.2%, 0.89 → 0.63). A parallel pattern emerged under balanced conditions. Removing the virtual node reduced AUROC by 3.4% (0.89 → 0.86), whereas eliminating JK caused a 7.9% decrease (0.89 → 0.82). Skip connections again proved critical: their removal led to a sharp 22.5% decline (0.89 → 0.69). The combined ablation of skip and JK resulted in a 16.9% drop (0.89 → 0.74), and exclusion of all components mirrored this effect (–22.5%, 0.89 → 0.69).

**Figure 4.**
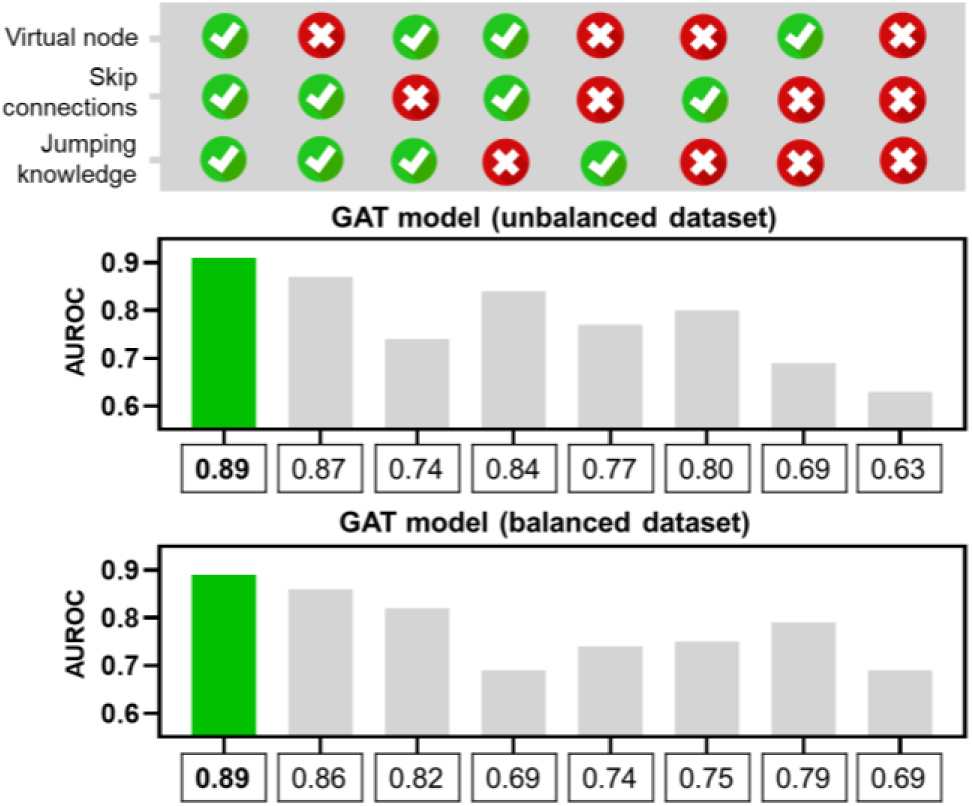
Ablation study of architectural components in holistic GAT models for antileishmanial activity prediction. (**a**) and (**b**) denote AUROC values for GAT models trained on the unbalanced and balanced datasets, respectively, evaluated across different combinations of architectural components.

### 2.5. Model explainability

To interpret model decisions, we prioritized counterfactual-based perturbation over attention weights, due to recent literature highlighting the unreliability of attention as a measure of feature importance.^43–45^ Besides assessing the model’s performance, it is often beneficial to examine the “black box” of the trained model to gain a deeper understanding of which substructures most influence antileishmanial activity. In this context, we implemented a counterfactual explainability approach using the GAT model trained on the balanced dataset, generating contribution maps by systematically perturbing atomic environments with isosteric and valence-based modifications and quantifying the resulting changes in predicted probabilities. This strategy aligns with recent advancements in model-agnostic counterfactual explainability for molecular property prediction, such as the framework proposed by Wellawatte et al.,^46^ which utilizes SELFIES-based molecular-level counterfactual generation to enhance interpretability in deep learning models.

As shown in Figure 5, the counterfactual maps revealed consistent positive contributions from well-established heterocyclic cores present in the literature series. In particular, 2,3-dihydroimidazo[2,1-*b*]oxazole (**a**),^47^ 6,7-dihydro-5H-imidazo[2,1-*b*][1,3]oxazine (**b**),^48–50^ benzoxaborole (**c**),^51^ imidazole (**d**),^52^ 2-benzil-1H-indole (**e**),^53^ quinoline (**f** and **g**),^54,55^ 1,3,5-triazine-2,4-diamine (**h**),^56^ cyanocarbamate (**i**),^57^ furane **(k**),^58^ tetrahydropyridine (**l** and **m**),^59^ and piperazine (**n**)^60^ moieties exhibited positive influence on predicted antileishmanial activity, underscoring their pharmacophoric relevance. In contrast, halogenated rings (**a-c**, **e**, **f**, **g**, **i**, and **j**) and unsubstituted phenyls (**d**, **l-n**) frequently contributed negatively within these scaffolds. These patterns indicate that the counterfactuality yielded chemically coherent interpretable maps that recapitulate established SARs—highlighting privileged heterocycles and penalizing fluorinated and unsubstituted phenyl motifs—and thereby support the selection and prioritization of key substructures for antileishmanial hit-to-lead optimization.

**Figure 5.**
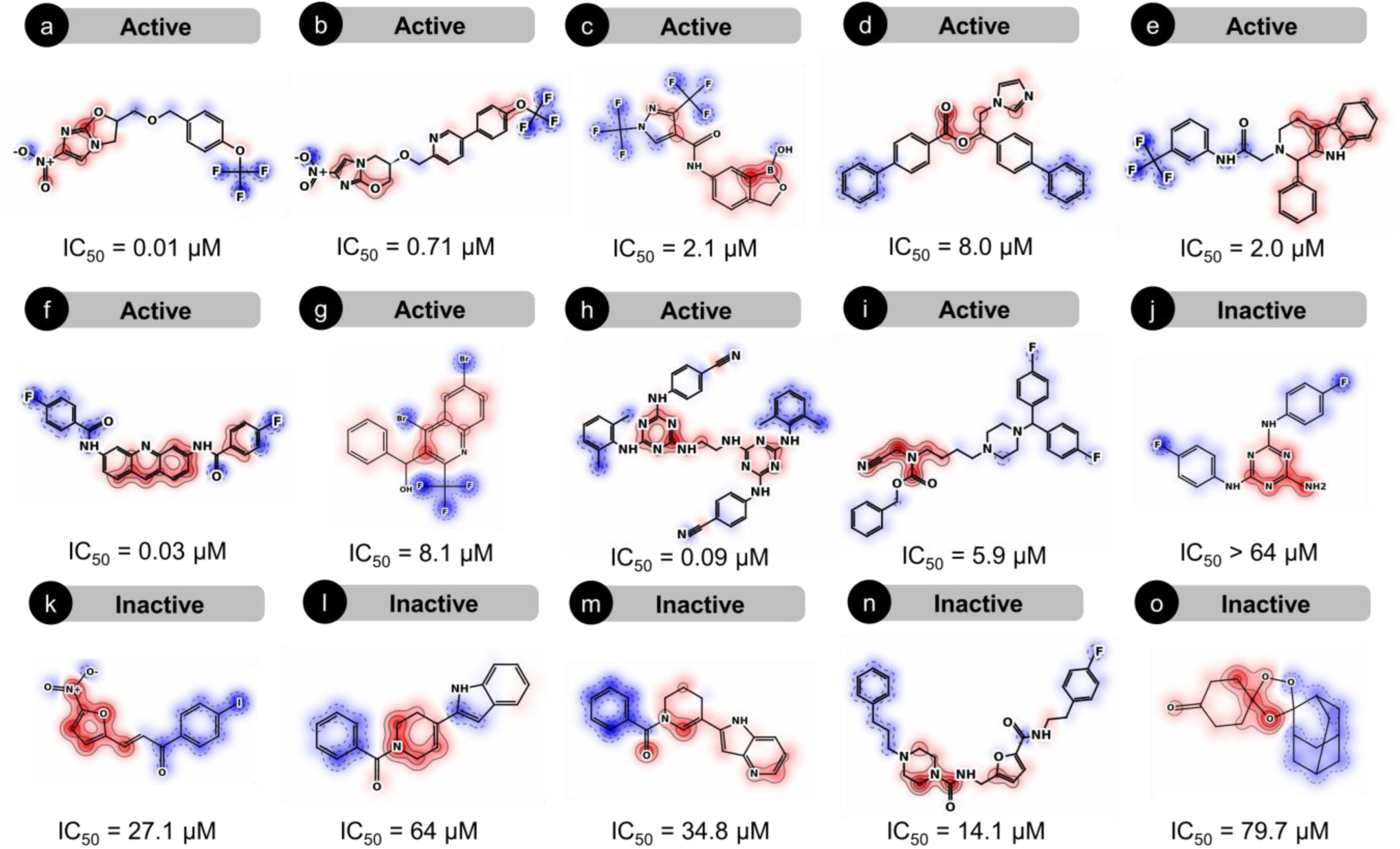
Counterfactual explainability of predicted antileishmanial activity. Red contours indicate regions contributing positively to antileishmanial activity, whereas blue contours indicate regions contributing negatively.

To further contextualize these findings, we explored attention-weight visualizations for the same set of molecules (Supplementary Figure S1) as an alternative strategy for model interpretability. However, these visualizations lacked clear patterns linking specific molecular substructures to predicted activity, limiting their practical interpretability in this setting. This limitation, consistent with recent literature questioning the reliability of attention weights as indicators of feature importance in GNNs,^45^ underscores the advantage of the counterfactual perturbation-based approach.

### 2.6. Virtual screening

A virtual screening workflow was designed to prioritize novel antileishmanial candidates from a commercially available library of 1.3 million compounds (Figure 6). Initially, the GAT model trained on the 10 µM activity threshold was applied as a primary filter, identifying approximately 20,000 compounds predicted as active with high confidence. Subsequently, the GAT model trained on the more stringent 1 µM activity threshold was employed as a secondary filter to refine the dataset, resulting in a subset of 5,000 compounds for downstream evaluation.

**Figure 6.**
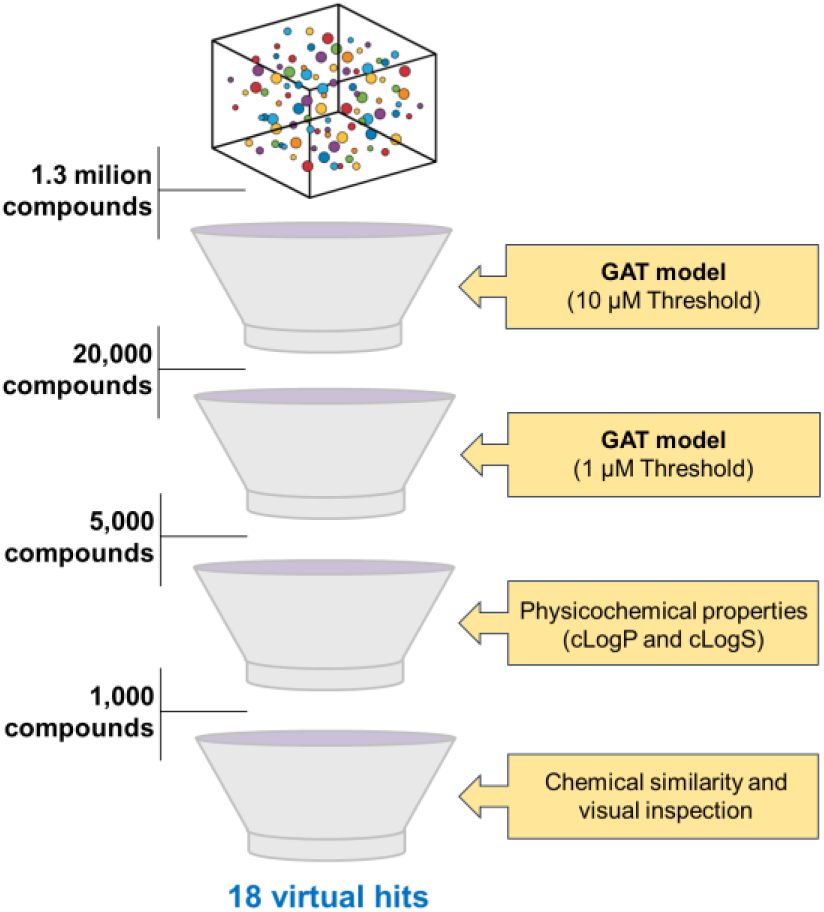
Virtual screening workflow for identifying novel antileishmanial compounds. The physicochemical filter was based on aqueous solubility (cLogS > -5.0) and lipophilicity (cLogP between 0.5 and 4.0). A structural novelty filter was applied using Tanimoto similarity with ECFP4 fingerprints to ensure that the putative hits had no prior experimental activity reported against *Leishmania* spp.

To ensure the drug-likeness and favorable pharmacokinetic properties, a physicochemical filter was applied based on aqueous solubility (cLogS > –5.0) and lipophilicity (cLogP between 0.5 and 4.0), narrowing the compound set to 1,000 structures. Finally, a structural novelty filter was implemented using Tanimoto similarity with ECFP4 fingerprints to exclude compounds with high similarity to known antileishmanial agents, ensuring the selection of structurally novel candidates. The nearest neighbors identified for these putative hits are provided in Table S4 for reference. Final selection incorporated medicinal chemistry expertise through visual inspection of the molecular structures to remove candidates with undesirable features. This multi-tiered screening cascade led to the identification of 18 putative hits, representing chemically diverse compounds suitable for further *in vitro* evaluation against *L. infantum*.

### 2.7. Experimental validation

The 18 putative hits were evaluated *in vitro* against promastigotes of *L. infantum* (Table 1). As a result, 11 compounds exhibited significant antileishmanial activity, with IC_50_ values ranging from 0.037 to 11.7 μM. To further characterize the prioritized hits, intracellular assays using THP-1-derived macrophages infected with *L. infantum* were conducted, followed by cytotoxicity assessments to determine the selectivity indexes.

**Table 1.**
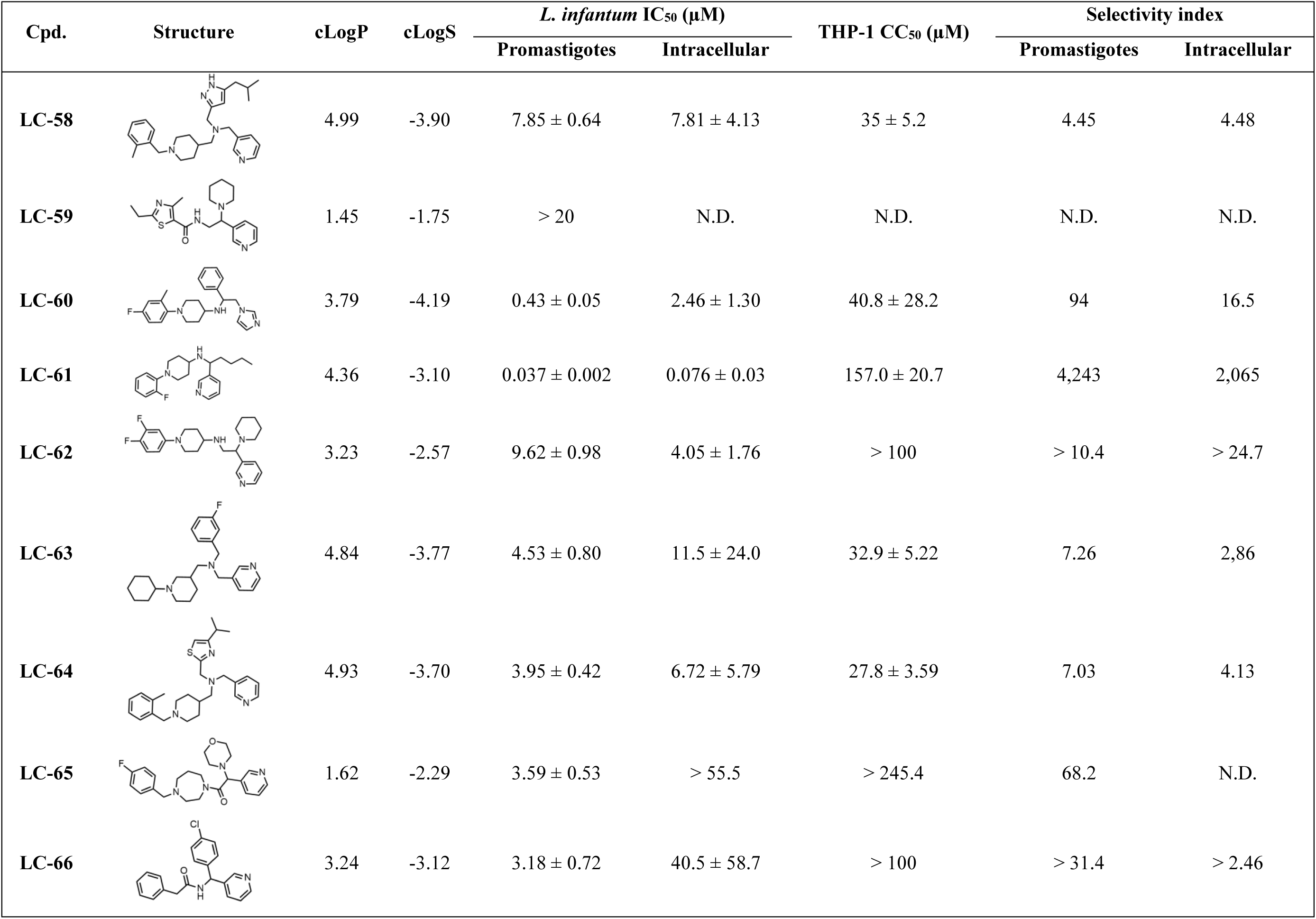
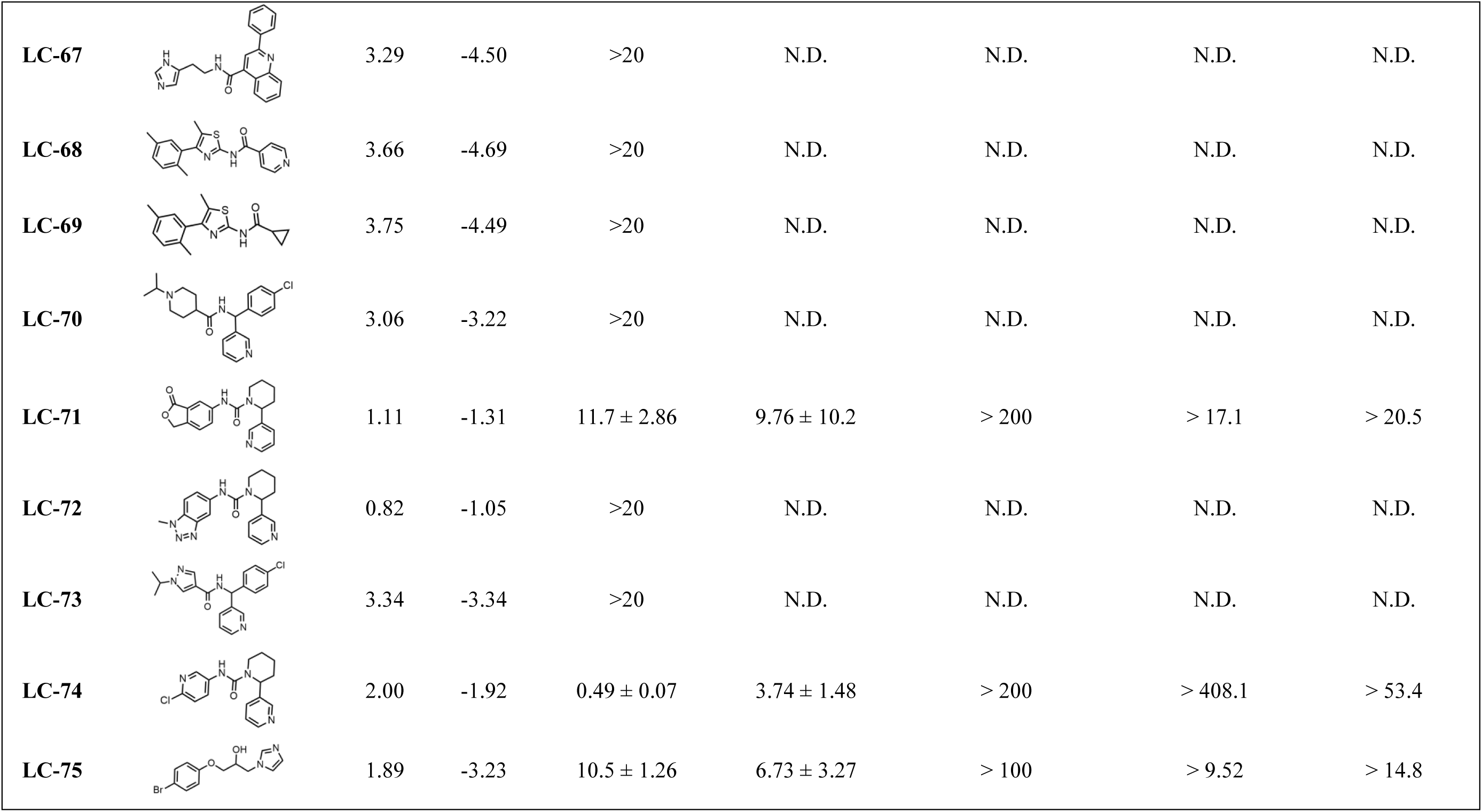
*In vitro* antileishmanial activity and selectivity of prioritized compounds against *L. infantum*.

| Cpd. | Structure | cLogP | cLogS | *L. infantum* IC<sub>50</sub> (μM) |  | THP-1 CC<sub>50</sub> (μM) | Selectivity index |  |
| --- | --- | --- | --- | --- | --- | --- | --- | --- |
|  |  |  |  | Promastigotes | Intracellular |  | Promastigotes | Intracellular |
| LC-58 |  | 4.99 | -3.90 | 7.85 ± 0.64 | 7.81 ± 4.13 | 35 ± 5.2 | 4.45 | 4.48 |
| LC-59 |  | 1.45 | -1.75 | > 20 | N.D. | N.D. | N.D. | N.D. |
| LC-60 |  | 3.79 | -4.19 | 0.43 ± 0.05 | 2.46 ± 1.30 | 40.8 ± 28.2 | 94 | 16.5 |
| LC-61 |  | 4.36 | -3.10 | 0.037 ± 0.002 | 0.076 ± 0.03 | 157.0 ± 20.7 | 4,243 | 2,065 |
| LC-62 |  | 3.23 | -2.57 | 9.62 ± 0.98 | 4.05 ± 1.76 | > 100 | > 10.4 | > 24.7 |
| LC-63 |  | 4.84 | -3.77 | 4.53 ± 0.80 | 11.5 ± 24.0 | 32.9 ± 5.22 | 7.26 | 2,86 |
| LC-64 |  | 4.93 | -3.70 | 3.95 ± 0.42 | 6.72 ± 5.79 | 27.8 ± 3.59 | 7.03 | 4.13 |
| LC-65 |  | 1.62 | -2.29 | 3.59 ± 0.53 | > 55.5 | > 245.4 | 68.2 | N.D. |
| LC-66 |  | 3.24 | -3.12 | 3.18 ± 0.72 | 40.5 ± 58.7 | > 100 | > 31.4 | > 2.46 |
| **LC-67** |  | 3.29 | -4.50 | >20 | N.D. | N.D. | N.D. | N.D. |
| **LC-68** |  | 3.66 | -4.69 | >20 | N.D. | N.D. | N.D. | N.D. |
| **LC-69** |  | 3.75 | -4.49 | >20 | N.D. | N.D. | N.D. | N.D. |
| **LC-70** |  | 3.06 | -3.22 | >20 | N.D. | N.D. | N.D. | N.D. |
| **LC-71** |  | 1.11 | -1.31 | 11.7 ± 2.86 | 9.76 ± 10.2 | > 200 | > 17.1 | > 20.5 |
| **LC-72** |  | 0.82 | -1.05 | >20 | N.D. | N.D. | N.D. | N.D. |
| **LC-73** |  | 3.34 | -3.34 | >20 | N.D. | N.D. | N.D. | N.D. |
| **LC-74** |  | 2.00 | -1.92 | 0.49 ± 0.07 | 3.74 ± 1.48 | > 200 | > 408.1 | > 53.4 |
| **LC-75** |  | 1.89 | -3.23 | 10.5 ± 1.26 | 6.73 ± 3.27 | > 100 | > 9.52 | > 14.8 |

As shown in Table 1, eight compounds maintained activity in the intracellular assay, with IC_50_ values ranging from 0.076 to 11.5 μM, underscoring the robustness of the screening cascade and the predictive power of our GNN framework. Four compounds (LC-58, LC-63, LC-64, and LC-66) exhibiting low intracellular potency (IC_50_ > 10 µM) and/or insufficient selectivity index (SI < 10) were deprioritized due to their limited potential for hit-to-lead optimization, as defined by established criteria for neglected tropical diseases.^38^ In contrast, six compounds (LC-60, LC-61, LC-62, LC-71, LC-74, and LC-75) demonstrated intracellular IC_50_ values below 10 µM while maintaining SIs > 17.

Among these validated hits, LC-60, LC-61, and LC-74 met our advancement criteria, displaying submicromolar to nanomolar activity and selectivity against intracellular amastigotes. LC-60 displayed submicromolar activity against intracellular amastigotes (IC_50_ = 2.46 µM) with low cytotoxicity in THP-1 cells (CC_50_ = 40.8 µM), yielding a favorable selectivity index (SI = 16.5). LC-74 likewise achieved submicromolar potency (IC_50_ = 3.74 µM) and low THP-1 cytotoxicity (CC_50_ > 200 µM), resulting in a similarly favorable SI (SI > 53.4). Notably, LC-61 emerged as the most promising hit, exhibiting potent nanomolar activity against both promastigotes (IC_50_ = 0.037 µM) and intracellular amastigotes (IC_50_ = 0.076 µM), combined with minimal cytotoxicity in THP-1 cells (CC_50_ = 157 µM) and an SI exceeding 2,000. Benchmarking against clinical comparators highlights LC-61’s advantage. Amphotericin B shows comparable potency against intracellular *L. infantum* amastigotes (IC_50_ = 0.14 µM) but exhibits high cytotoxicity (CC_50_ = 1.2 µM), yielding a low selectivity index (SI = 8.6).^61^ Similarly, miltefosine demonstrated lower potency (IC_50_ = 0.37 µM) and moderate cytotoxicity (CC_50_ = 51.1 µM, SI = 138).^61^

### 2.8. Early ADME profiling of hits

To prioritize candidates for hit-to-lead optimization, *in vitro* ADME profiles were determined for compounds LC-60, LC-61, and LC-74, the most potent hits against *L. infantum* amastigotes (IC_50_ < 5 μM). Comprehensive experimental details and analytical data are provided in the Supporting Information (Files S1–S5). As shown in Table 2, all three hits showed high aqueous solubility at acidic and physiological pHs (≈28–33 µg/mL). At pH 2.0, LC-60 and LC-61 exceeded 28 µg/mL (>32.21 and >28.35 µg/mL, respectively), while LC-74 reached 27.58 µg/mL; at pH 7.4, LC-74 was the most soluble (32.81 µg/mL), followed by LC-60 (>30.03 µg/mL) and LC-61 (>28.18 µg/mL). Experimentally measured distribution coefficients (eLogD) were within a drug-like range: LC-60 and LC-61 showed values near the Lipinski Rule-of-Five threshold (4.45 and 4.07, respectively), whereas LC-74 was less lipophilic (2.39). Consistent with the observed eLogD values, parallel artificial membrane permeability assays (PAMPA) showed high apparent passive permeability for all three compounds (4.86–10.85 × 10^-6^ cm/s): LC-74 was highest (10.85 × 10^-6^ cm/s), followed by LC-60 (6.36 × 10^-6^ cm/s) and LC-61 (4.86 × 10^-6^ cm/s).

**Table 2.**
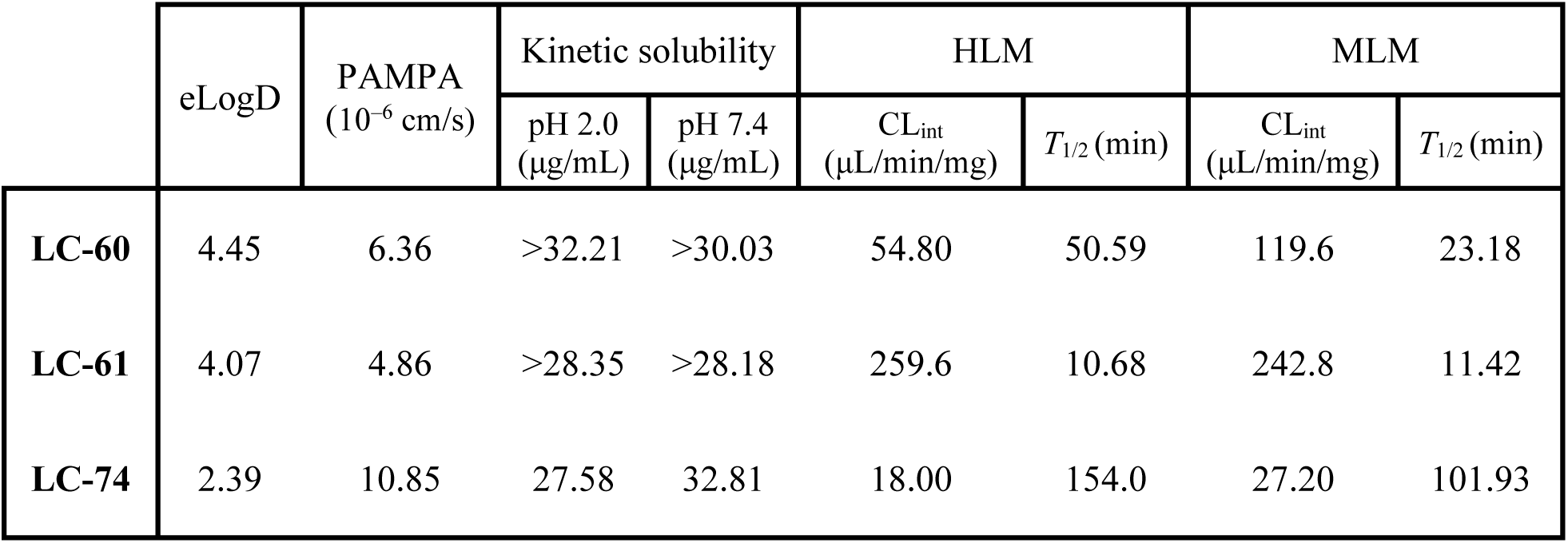
Summary of *in vitro* ADME properties for top compounds.

Microsomal stability assays revealed distinct metabolic profiles among the three hits (Table 2). LC-60 exhibited moderate intrinsic clearance in human liver microsomes (HLM, CL_int_ = 54.80 µL/min/mg) with shorter half-life (*T*_1/2_ = 50.59 min), while clearance in mouse liver microsomes (MLM) was lower (CL_int_ = 119.6 µL/min/mg; *T*_1/2_ = 23.18 min), suggesting species-dependent turnover. LC-74 showed the highest microsomal stability across both systems, with low clearance values (MLM CL_int_ = 27.20 µL/min/mg; HLM CL_int_ = 18.0 µL/min/mg) and adequate half-lives (101.93 and 154.0 min, respectively), consistent with its lower lipophilicity (eLogD = 2.39). In contrast, LC-61 exhibited a pronounced metabolic turnover, characterized by higher intrinsic clearance in both species (MLM CL_int_ = 242.8 µL/min/mg; HLM CL_int_ = 259.6 µL/min/mg) and correspondingly short half-lives (MLM *T*_1/2_ = 11.42 min; HLM *T*_1/2_ = 10.68 min). Notably, the elevated clearance of LC-61 occurred despite its balanced lipophilicity (eLogD = 4.07), suggesting that rapid turnover stems from structural liabilities rather than global physicochemical properties.

The presence of an unprotected aromatic ring likely renders the molecule vulnerable to hydroxylation catalyzed by cytochrome P450 isoforms, consistent with the higher clearance observed in human microsomes. Such metabolic instability could be mitigated through targeted electronic modulation (e.g., selective fluorination or methoxylation) to shield the reactive aromatic site while preserving potency and physicochemical balance. Despite its moderate metabolic liability, LC-61 remains the most promising hit, combining nanomolar potency, favorable permeability, and an overall drug-like profile that strongly supports its advancement into hit-to-lead optimization. In addition, LC-61 demonstrates favorable synthetic accessibility, with a SwissADME^62^ Synthetic Accessibility Score (SAscore) of 3.18, indicating that the compound can be obtained through feasible synthetic routes and supporting its potential for further lead optimization.

### 2.9. Target fishing

To elucidate the potential mechanism of action of LC-61, a target fishing analysis was implemented through a multi-step *in silico* approach. Initially, the pyridine moiety highlighted in the contribution maps of LC-61 and its nearest neighbors (Figure 7a) was used for substructure-based searches in the ChEMBL database and scientific literature. This approach supported the identification of potential targets associated with these moieties. Based on this rationale, *L. infantum* sterol C4-methyl oxidase (CYP5122A1)^63,64^ and sterol 14α-demethylase (CYP51)^65,66^ were prioritized for further evaluation due to their documented tractability as targets for pyridine-based inhibitors. These cytochrome P450 enzymes catalyze sequential oxidative steps in the ergosterol biosynthesis pathway in *Leishmania*, with CYP5122A1 forming C4-oxidation metabolites and CYP51 mediating sterol 14α-demethylation.^63,64^

**Figure 7.**
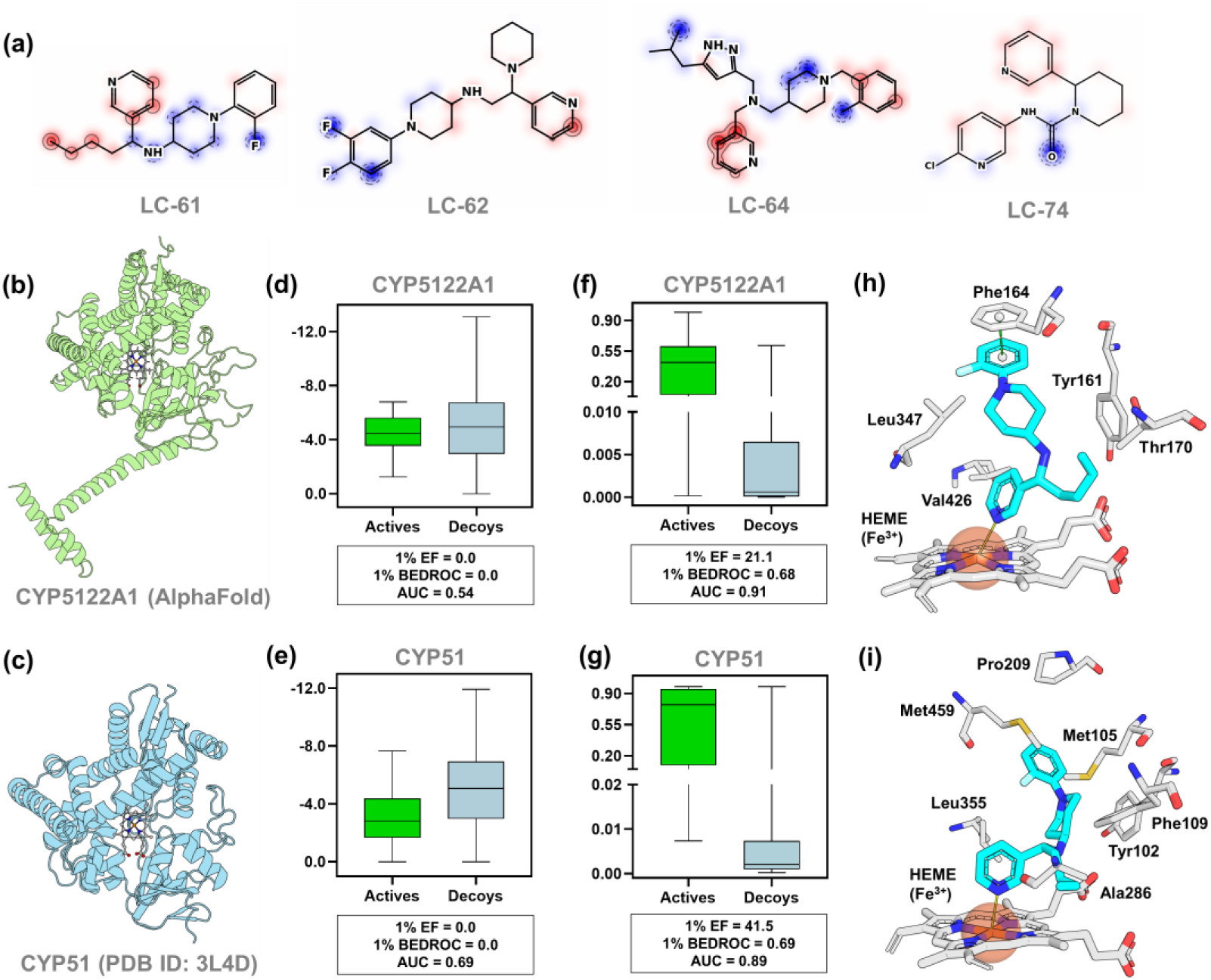
Target fishing and molecular docking of LC-61. (**a**) Counterfactual explainability maps for LC-61 and its neighbors, evidencing positive contributions concentrated on the pyridine ring. Red contours indicate regions contributing positively to antileishmanial activity, whereas blue contours indicate regions contributing negatively. (**b**) Predicted 3D structure of *L. infantum* CYP5122A1 and (**c**) X-ray structure of *L. infantum* CYP51 (PDB ID: 3L4D), respectively. Docking scores and enrichment metrics for (**d**) CYP5122A1 and (**e**) CYP51 obtained using a benchmark dataset of known actives and decoys. Bayesian rescoring and enrichment metrics for (**f**) CYP5122A1 and (**g**) CYP51 obtained using a benchmark dataset of known actives and decoys. Predicted binding poses of LC-61 in the active sites of (**h**) CYP5122A1 and (**i**) CYP51, highlighting key interactions with active site residues and HEME.

Subsequently, molecular docking simulations were performed against the crystal structure of *L. infantum* CYP51^67^ (Figure 7b) and the AlphaFold-predicted structure of *L. infantum* CYP5122A1 (Figure 7c and File S2). However, initial analysis of docking scores (GlideScore function) using a benchmark dataset of known actives and decoys yielded poor enrichment performance, with CYP5122A1 achieving an AUROC of 0.54 and both Boltzmann-Enhanced Discrimination of ROC (BEDROC_₁%_) and Enrichment Factor (EF_₁%_) values of 0.0 (Figure 7d). At the same time, CYP51 showed an AUROC of 0.69 with BEDROC₁% and EF₁% similarly at 0.0 (Figure 7e). These outcomes demonstrate that GlideScore failed to discriminate actives from decoys for either target, with AUROCs near random expectation and no early enrichment, thereby lacking target-fishing utility for reliable target assignment.

To enhance the reliability of the docking protocol, a Bayesian rescoring function was developed using raw energy terms and scoring descriptors extracted from the docking outputs. This approach substantially improved predictive performance across all metrics, with CYP5122A1 reaching an AUROC of 0.91, BEDROC_1%_ of 0.68, and EF_1%_ of 21.1 (Figure 7f), and CYP51 achieving an AUROC of 0.89, BEDROC_1%_ of 0.69, and EF_1%_ of 41.5 (Figure 7g). While initial docking yielded limited enrichment, Bayesian rescoring significantly improved discrimination, underscoring the limitations of raw docking scores for novel scaffolds. These results confirm the effectiveness of the Bayesian rescoring strategy in enhancing the docking protocol’s ability to distinguish actives from decoys.

Following validation of the docking protocol, LC-61 emerged as the only candidate predicted to inhibit both CYP5122A1 and CYP51. In CYP5122A1 (Figure 7h), LC-61 adopted a stable binding pose within the active site, with the fluorobenzene ring forming a π–π stacking interaction with Phe164, while the butyl substituent engaged in hydrophobic contacts with Tyr161 and Thr170, contributing to the stabilization of the ligand within the binding pocket. Notably, the pyridine nitrogen was oriented for axial coordination with HEME iron, a key interaction commonly exploited by CYP inhibitors.

In CYP51 (Figure 7i), LC-61 also displayed a favorable binding orientation, with fluorobenzene engaging hydrophobic contacts with Met105, Pro209, and Met459. In addition, the pyridine nitrogen was positioned for axial coordination with the HEME iron, mirroring the key interaction observed in CYP5122A1, while the butyl substituent engaged in hydrophobic contacts with Ala286, Phe109, and Tyr102. These findings are consistent with recent work demonstrating that dual inhibition of CYP5122A1 and CYP51 is required for optimal antileishmanial activity, as inhibition of CYP51 alone results in limited efficacy due to compensatory pathways in ergosterol biosynthesis.^68^ Notably, LC-61 was the most potent compound identified in this study, supporting the notion that concurrent blockade of C4-methyl oxidation and C14-demethylation enhances antileishmanial efficacy. Importantly, the scaffold of LC-61 offers multiple opportunities for derivatization, which could enable fine-tuning of potency and metabolic stability.

## 3. CONCLUSIONS

In this study, we hypothesized that integrating chemical semantics and long-range structural context into GNNs would improve the prediction of antileishmanial activity and facilitate the identification of novel antileishmanial compounds. This architecture achieved substantial predictive gains over conventional GNNs and successfully integrated counterfactual explainability, enabling atom-level interpretation of model predictions and rational prioritization of structural motifs driving antileishmanial activity. The application of this framework to a large and chemically diverse library of 1.3 million compounds yielded 18 putative hits with a 50% hit rate during *in vitro* experimental validation, underscoring its practical utility in accelerating antileishmanial hit discovery. Among these, LC-61 emerged as a standout hit-to-lead candidate, featuring a chemically novel scaffold with nanomolar potency against intracellular amastigotes (IC_50_ = 0.076 µM), low cytotoxicity (CC_50_ = 157 µM), and a highly favorable SI (> 2000). Its balanced kinetic solubility (> 28 µg/mL at pH 2.0–7.4), lipophilicity (eLogD = 4.07), and high passive permeability (PAMPA = 4.86×10^-6^ cm/s) further support its advancement despite moderate microsomal turnover, which can be mitigated through targeted electronic modifications in aromatic ring. These findings highlight LC-61 as a validated and pharmacokinetically tractable hit ready for hit-to-lead optimization and demonstrate the broader potential of our GNN framework to streamline discovery of novel antileishmanial compounds.

## 4. EXPERIMENTAL SECTION

### 4.1. Data collection and curation

Compounds with IC_50_ data obtained after 72 hours of exposure against intracellular amastigotes of *Leishmania infantum* (CHEMBL612848) were retrieved from the ChEMBL database.^37^ Subsequently, samples were labeled as active or inactive using activity thresholds of 1 μM and 10 μM, providing clear class delineation for subsequent predictive modeling analyses. Chemical structures in simplified molecular-input line-entry system (SMILES) format and corresponding bioactivity data were meticulously curated following the guidelines proposed by Fourches et al.^69,70^ Curation steps included the normalization of nitro groups and aromatic systems, removal of salts, mixtures, polymers, and organometallic compounds, as well as standardization of tautomeric forms to ensure consistency across the dataset. For duplicate entries with discordant activities, all corresponding records were excluded, while duplicates with concordant activities were consolidated to a single representative entry.

### 4.2. Data description

Chemical diversity within the dataset was assessed by calculating pairwise molecular similarity distributions using ECFP4 fingerprints with Tanimoto, Dice, and Asymmetric similarity metrics. Molecular fingerprints were generated using the RDKit framework v.2024.4.5, and similar distributions were visualized using kernel density estimates. In parallel, physicochemical profiles were characterized using cLogP, MW, number of rotatable bonds, and tPSA computed for each compound using RDKit descriptor functions. Subsequently, scatter plot matrices and marginal density plots were generated to visualize the distribution of these physicochemical descriptors across actives and inactives.

### 4.3. Holistic GNNs

#### 4.3.1. Split of datasets

The curated datasets were randomly divided into training, validation, and test sets in an 8:1:1 ratio. The training set was used to build the model, the validation set for hyperparameter tuning, and the test set for model evaluation.

#### 4.3.2. Feature representation

Molecular graphs were generated from SMILES using RDKit, with atoms represented as nodes and bonds as edges. Atom-level features included one-hot encoding of atomic number, degree, formal charge, hybridization state, and aromaticity. Bond-level features were encoded using one-hot representations of bond type, conjugation status, ring membership, and stereochemistry, including chirality. Feature vectors were built as tensors to provide structured graph representations during model training.

#### 4.3.3. Architectural innovations

First, skip connections enable the models to combine low-level and high-level features, facilitating better training dynamics and expressiveness as follows:

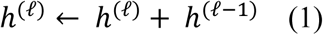

Where ℎ^(ℓ−1)^, ℎ^(ℓ)^ ∈ ℝ^𝑑^ are input and output features at layer ℓ. This additive transform acts as identity initialization, preserving gradient flow. For dimension mismatches, linear projections align features before addition.

In parallel, a virtual node was incorporated. This node is initialized with a learnable embedding by 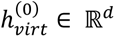 and interacts with all atomic nodes by contributing additively to their representations at each message-passing layer. For a given graph 𝐺 = (𝑉, 𝐸), the representation of each atom 𝑣 ∈ 𝑉 at layer ℓ is modified as follows:

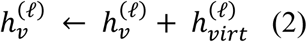

where 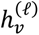 is the atom’s representation at layer ℓ, and 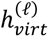 is the virtual node’s representation at the same layer. This additive interaction ensures that global molecular information is dynamically propagated to all atomic nodes. After message passing, the virtual node embedding is updated by aggregating the current node states across graph:

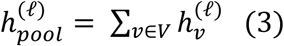

where 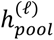 is the graph-level summary at layer ℓ, computed as permutation-invariant sum over all atomic representations 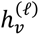. The pooled representation is then transformed through a learnable nonlinear function 𝑀𝐿𝑃^(ℓ)^, typically consisting of two linear layers with intermediate activation and normalization, to produce a virtual node embedding 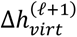:

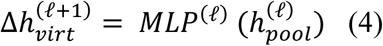

This ensures the virtual node serves as a contextual relay, iteratively refining global semantics while retaining memory of prior states. Finally, the virtual node embedding is updated by residual addition:

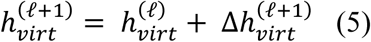

Where the virtual node embedding from the previous layer 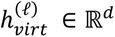 is incremented by the newly computed update 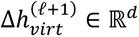. This residual update scheme allows the virtual node to dynamically absorb and redistribute information, functioning as a central relay for integrating subgraph semantics and contextualizing local features.

For the Jumping Knowledge, let H^(0)^, H^(1)^, . . ., H^(𝐿)^ denote the sequence of node embeddings obtained at each of the *L* message-passing layers. JK aggregation builds a unified node representation through feature concatenation:

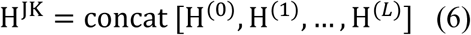

This final embedding H^JK^ ∈ ℝ^|𝑉|^ ^x^ ^𝑑^_JK_, where 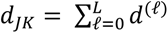, captures multiscale structure features by aggregating both shallow and deep node contexts. Subsequently, a graph-level embedding ℎ_𝐺_ ∈ ℝ ^𝑑^_JK_ is obtained via pooling function over node embeddings as follows:

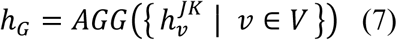

Depending on the architecture and task requirements, AGG denotes a permutation-invariant function such as summation, averaging, or attention-weighted pooling. This strategy ensures that the model captures the hierarchical nature of chemical graphs, from local functional groups to extended rings and substructures, thereby enhancing its ability to generalize across complex molecules.

#### 4.3.4. Model training and evaluation

Holistic GNNs were implemented and trained using PyTorch v.2.5.1 and PyTorch Geometric v.2.6.1 on an NVIDIA TITAN Xp GPU. On-the-fly batches with sizes of 64 and 48 were used for the unbalanced and balanced datasets, respectively. Models were trained for a maximum of 2,000 epochs using binary cross-entropy (BCE) loss, and early stopping was applied to prevent overfitting. Training terminates when no improvement in validation loss is observed within the adaptive five-epoch window; otherwise, the best validation loss checkpoint is refreshed. Regularization strategies, including gradient clipping and learning rate scheduling (cosine annealing with warm-up), were employed to improve generalization. Pruning strategies were applied during optimization to terminate unpromising trials early. Hyperparameter optimization was conducted using Optuna v4.2.0, with 50 trials per architecture to explore the search space systematically. Model performance was evaluated using ACC, recall, SP, and AUROC, with G-mean additionally considered to address the imbalanced nature of the datasets. Finally, embedding vectors were saved during training and projected for visualization using t-SNE and kernel density estimation (KDE) to monitor the organization of the latent space across epochs.

#### 4.3.5. Model calibration

A probability calibration procedure was implemented using a threshold-moving strategy ^71^ to refine decision boundaries derived from raw predicted probabilities. A threshold sweep (0.0–1.0, increment 0.01) was performed on the training and validation sets, selecting thresholds that maximized the G-mean. The calibrated thresholds were then applied to the test set to reclassify compound predictions, improving the balance between sensitivity and specificity while preserving the AUROC.

#### 4.3.6. Model explainability

A counterfactual approach was developed to generate atom-level contribution maps. Initially, stochastic perturbations were applied to molecular graphs using rule-based isosteric and valence-preserving substitutions, quantifying positive and negative contributions relative to the baseline prediction of the unperturbed molecule. For each atom 𝑖, the importance score was defined as:

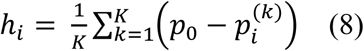

where 𝑝_0_ is the predicted probability for the unperturbed graph, 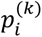 is the predicted probability after *k*-th perturbation of atom 𝑖, and 𝐾 is the number of perturbation samples. Graph-based clustering was then performed using connected components to group structurally connected atoms with similar contribution signals, with intra-cluster enhancement applied as:

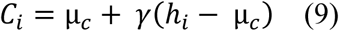

where 𝐶*_i_* is the cluster-enhanced contribution score for atom 𝑖, 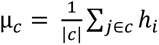 is the mean contribution 𝑐, and 𝛾 > 0 is the cluster-enhacement factor. Laplacian smoothing was subsequently applied within each cluster to reduce noise while preserving local structure as follows:

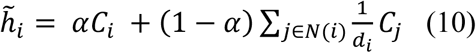

where 𝐶_𝑗_ is the cluster-enhanced score of each neighbor 𝑗 ∈ 𝑁(𝑖), whereas 𝛼 ∈ [0, 1] balances self-vs-neighbor contributions, 𝑁(𝑖) is the set of neighbors of atom 𝑖, and 𝑑_𝑖_ = |𝑁(𝑖)| its degree. Finally, the 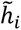 values are normalized (zero mean, unit variance) and projected onto 2D chemical structures via RDKit’s similarity maps for visualization. Additionally, atom-level attention weights from attention-based GNNs were extracted and smoothed using the same clustering and Laplacian strategies, providing a complementary visualization of substructural relevance.

### 4.4. Target fishing

#### 4.4.1. Substructure search

Initially, the pyridine motif was prioritized based on the LC-61 counterfactual map and encoded as SMARTS pattern. SMARTS-based substructure research in ChEMBL and a targeted literature review were conducted to collect bioactivity data for *Leishmania* proteins whose ligands matched the encoded motifs. *L. infantum* target candidates were then limited to enzymes with reported roles in sterol biosynthesis, identifying CYP51 and CYP5122A1 for subsequent *in silico* evaluation.

#### 4.4.2. Protein preparation

The X-ray structure of CYP51 (PDB ID: 3L4D) and the AlphaFold2-predicted CYP5122A1 model (UniProt ID: A4I2K5) were processed using Protein Preparation Wizard in Maestro workspace v.9.3 (Schrödinger, LCC, New York, 2012). In this step, hydrogen atoms were added to the proteins, and bond orders and formal charges were adjusted accordingly. Further, the Epik^72^ was employed to predict the protonation states (p*K*_a_) of polar amino acids at pH = 7.4 ± 0.5, whereas the PROPKA v.3.1^73,74^ was used to optimize the hydrogen orientations. Subsequently, loop regions in the CYP5122A1 model were rebuilt using Prime’s loop refinement (Schrödinger, LCC, New York, 2012). This procedure samples alternative backbone conformations and optimizes loop geometry under the OPLS2005 force field,^75^ ensuring a more accurate active-site architecture for subsequent docking.

#### 4.4.3. Ligand preparation

A series of 26 CYP5122A1 and 37 CYP51 inhibitors (IC_50_ ≤ 10 µM) was compiled from the literature. Then, decoys were generated for each inhibitor via the LUDe server, yielding a total of 1,300 decoys for CYP5122A1 and 1,850 decoys for CYP51. All actives, decoys, and prioritized hits were processed with LigPrep 2.5 to generate low-energy conformers and predict protonation states at pH 7.4 ± 0.5.

#### 4.4.4. Molecular docking

Grid boxes were established into *x*, *y*, and *z* coordinates of CYP5122A1 (37.64 x -25.58 x -49.06) and CYP51 (37.64 x -25.68 x -49.25) using the receptor grid generation panel of Glide v.5.8. Then, molecular docking calculations were performed on the Maestro workspace Glide v.5.8,^76,77^ employing extra precision (XP) mode. The poses were scored using GlideScore and further optimized using the OPLS2005 force field.^75^ Docking energy terms and scores were exported and used to train a Naive Bayes classifier in KNIME v.5.2.5 with five-fold cross-validation. The performance of the docking protocol and Bayesian rescoring model was assessed using EF, BEDROC, and AUROC metrics.

### 4.5. Experimental validation

#### 4.5.1. Chemicals

Compounds were purchased from ChemBridge (San Diego, CA, USA), dissolved in 100% dimethylsulfoxide (DMSO) to achieve a stock concentration of 20 mM, and stored at -20 °C. The chemical structures of all compounds were verified using proton nuclear magnetic resonance (^1^H NMR) spectroscopy or liquid chromatography-mass spectrometry (LC-MS) analysis, which included evaporative light scattering and ultraviolet detectors. This analysis confirmed that all compounds had a minimum purity of 95% (spectra of the compounds are provided in the Supporting Information).

#### 4.5.2. Parasite and cell culture

*L. infantum* promastigotes were grown at 26°C in Schneider medium (Sigma), supplemented with 20% heat-inactivated fetal bovine serum (Sigma) and 1% penicillin-streptomycin (Sigma). To ensure maintenance of virulence, the parasites were serially passed through BALB/c mice. THP-1 cells were cultured in RPMI-1640 medium (Sigma-Aldrich), supplemented with 10% heat-inactivated SFB (Sigma) and 1% penicillin-streptomycin (Life Technologies), maintained at a density of 10⁶ cells/mL, and incubated at 37 °C in a 5% CO_2_ atmosphere. For differentiation, THP-1 cells were seeded in 96-well culture plates at a concentration of 1×10⁴ cells/well and treated with 100 ng/mL of 4α-phorbol 12-myristate 13-acetate (PMA, Sigma-Aldrich) and incubated for 48 hours at 37 °C and 5% CO_2_. Cell viability was previously assessed using the Trypan Blue exclusion method. After stimulation with PMA, the cells exhibited characteristics of macrophages, including morphological changes and adherence to the culture plates. Differentiated THP-1 cells were washed with non-supplemented RPMI-1640 and given fresh medium without PMA. The cells were rested for 24 hours before further treatment.

#### 4.5.3. Promastigote assay

To evaluate the effect of prioritized compounds on parasite viability, 4 × 10^6^ promastigotes/mL were incubated in the presence of a series of 11 points from 200–0.195 or 50–0.04 μM plus control (0.1% DMSO) for 48 h at 28 °C to a final volume of 200 μL in a 96-well microplate. Next, 20 μL of resazurin solution (0.39 mM) was added to each well and incubated for 4 h, and then fluorescence was recorded (570 nm_ex_/595 nm_em_) on a microplate reader SpectraMax M5 (Molecular Devices, Sunnyvale, CA). The IC_50_ from three independent experiments was calculated from the dose–response curve fitted in a non-linear regression using a four-parameter logistic (4PL) model in GraphPad Prism v.10.

#### 4.5.4. Intracellular assay

THP-1 cells differentiated in 96-well culture plates were infected with promastigotes in the stationary phase at a ratio of 30 parasites: 1 cell for 4 hours at 37 °C in a final volume of 100 µL. The non-internalized parasites were then removed by washing with RPMI-1640 medium. The compounds were diluted at concentrations of 200-0.7 µM in 100 µL of culture medium per well for serial dilution. Subsequently, 100 µL of the diluted compounds were added to the infected THP-1 cells and incubated for 48 hours. The wells were stained with panotype, and the number of intracellular parasites per 100 macrophages was counted using light microscopy. The infection rate was determined in relation to the mean of the control wells, and the IC_50_ calculation was performed using a 4PL model in GraphPad Prism v.10.

#### 4.5.5. Cytotoxicity assay

THP-1 cells were cultured in 96-well plates at a density of 1 × 10⁴ cells per well in 100 µL of RPMI 1640 medium supplemented with 10% heat-inactivated fetal bovine serum (FBS) and 25 mM HEPES (pH 7.4). Cultures were maintained at 37 °C in a humidified atmosphere containing 5% CO₂. After 24 hours of incubation, the test compounds were prepared in serial dilutions in complete medium, starting at a concentration of 200 µM. Subsequently, 100 µL of each diluted compound solution was added to the corresponding wells containing the cells. Following a 72-hour incubation period, resazurin solution (Sigma^®^) was added to each well at a final concentration of 20 µM, and plates were incubated for an additional 4 hours under the same conditions. Fluorescence was measured using a SpectraMax M5 microplate reader (Molecular Devices, Sunnyvale, CA, USA) with an excitation wavelength of 570 nm and an emission wavelength of 595 nm. The cytotoxic concentration that reduced cell viability by 50% (CC_50_) was determined based on data from two independent experiments and calculated using non-linear regression analysis, as previously described for IC_50_ determination.

#### 4.5.6. In vitro ADME characterization

ADME profiling was conducted using liquid chromatography coupled with tandem mass spectrometry (LC–MS/MS). The chromatographic separation was carried out on a Prominence UFLC system (Shimadzu Corporation, Kyoto, Japan), connected to an LCMS-8045 triple quadrupole mass spectrometer (Shimadzu Corporation, Kyoto, Japan) with an electrospray ionization.

##### 4.5.7.1. Kinetic solubility

Kinetic solubility of the test compounds was assessed by preparing 10 mM stock solutions in DMSO, which were then dispensed into two 96-well incubation plates, each in duplicate. For each well, either PBS at pH 7.4 or 2.0 was added to reach a final compound concentration of 250 μM, keeping the DMSO content below 2.5%. The plates were sealed and shaken (200 rpm) at 25 °C for 24 h to allow for equilibrium solubility to be reached. The precipitates on the incubation plate were removed by centrifugation at 3000 rpm for 15 min at 25 °C, and the supernatant fractions were quantified by LC-MS/MS. An intermediate standard solution diluted to 0.5 mM in a 1:1 solution of acetonitrile and water. A calibration curve was prepared for each of the test compounds and controls by diluting several times the intermediate standard solution to reach the desired concentrations of 50, 40, 20, 2, and 1 μM. The resulting equation for the calibration (*y = mx + b*) was used to calculate the actual concentrations present in the test samples. The chromatogram for analysis was achieved on a Supelco Ascentis Express C18 column (3 cm × 2.1 mm, 5 μm particle size) using water with 0.05% formic acid (A) and acetonitrile with 0.05% formic acid (B) as mobile phase. The mobile phase was eluted in binary gradient mode, and the gradient was as follows: 0 min: 98% A; 1.2 min: 2% A; 2.0 min: 2% A; re-equilibration time: 0.6 min, 98% A. The total run time was 2 min per sample, with an injection volume of 5 μL and a flow rate of 0.6 mL/min.

##### 4.5.7.2. Experimental determination of distribution coefficient (eLogD)

The eLogD was carried out using a chromatographic method based on the analyte retention times within a stationary phase. Separation was performed on a Supelco Ascentis Express RP Amide high-performance liquid chromatography (HPLC) column (5 cm × 2.1 mm, 2.7 μm particle size), utilizing a binary mobile phase system composed of 5% methanol in 10 mM ammonium acetate buffer at pH 7.4 (designated as solvent A), and pure methanol (solvent B). The gradient program was as follows: initial composition at 95% A; shifted to 100% A at 0.3 min; reduced to 0% A by 5.2 min; maintained at 0% A until 5.6 min; returned to 100% A at 5.8 min; and held until the end of the 7 min run. The injection volume for each sample was 5 μL. Test compounds were diluted to a concentration of 1.0 mg/mL in a 1:1 mixture of mobile phases A and B, containing an internal standard at 200 nM. Final DMSO concentration was kept below 2%. To determine compound lipophilicity, each test molecule was injected individually along with a panel of eight reference drugs with known eLogD_7.4_ values ranging from −1.86 to 6.10. These standards included: acyclovir (−1.86), atenolol (0.16), antipyrine (0.38), fluconazole (0.50), prednisone (1.46), ketoconazole (3.83), tolnaftate (5.40), and amiodarone (6.10). A calibration curve was generated by plotting the retention times of these standards against their corresponding eLogD values. The linear regression equation (*y* = *mx* + *b*) obtained from this curve was then used to calculate the eLogD of each test compound.

##### 4.5.7.3. Parallel artificial membrane permeability assays

To evaluate the effective permeability (Pe) of the compounds, a pre-coated 96-well BD Gentest™ PAMPA plate (Corning Gentest #353015) is used. Each well is divided into two chambers, donor and acceptor, separated by a triple-layer phospholipid membrane constructed on a porous filter. Solutions of the compounds are prepared by diluting their respective stock solutions (10 mM in DMSO) in phosphate-buffered saline (PBS) at pH 6.5, yielding a final concentration of 10 µM (<1% DMSO, v/v). The solutions are then added to the donor portion of the plate (300 µL/well) in triplicate, while the acceptor portion received only PBS pH 7.4 (200 µL/well). The donor and acceptor portions are subsequently assembled, and the system is incubated for 5 hours at 37°C and 100 rpm. Samples of the initial donor solution (T_zero_) are collected and transferred to an analysis plate (10 µL), followed by the addition of stop solution (300 µL) [10% ultrapure water (type 1) and 90% methanol (HPLC grade ≥99.9%):acetonitrile (HPLC grade ≥99.9%) (50:50) + 50 nM internal standard] and PBS buffer pH 6.5 (60 µL). After the incubation period, samples are collected from the donor (10 µL) and acceptor (80 µL) wells and added to the analysis plate containing stop solution (300 µL) and PBS buffer pH 6.5 (60 µL). Final compound concentrations in the donor, acceptor, and T_zero_ wells are quantified by LC-MS/MS. LC-MS/MS. The chromatogram for analysis was achieved on Supelco Ascentis express C18 column (3cm x 2.1mm, 5µM). The mobile phases consisted of water + 0.1% formic acid (A) and acetonitrile + 0.1% formic acid (B). Mobile phase was eluted in binary gradient mode, and the gradient was as follows: 0 min., 95% A; 0.05 mins. 95% A; 0.3 mins: 2% A; 0.7 mins: 2% A; 0.8 mins: 95% A; 1.15 mins 95% A; 2.0 mins 95% A. The run time was 2 minutes, and the sample injection volume was 10 µL and flow rate of 0.7 mL/min. The results were used to calculate an effective permeability (Pe) value. All PAMPA assay was performed in triplicates.

##### 4.5.7.4. Human and mouse liver microsomal stability assay

Metabolic stability of the test compounds was assessed using pooled human liver microsomes (20 mg/mL, GIBCO) and CD1 mouse liver microsomes (20 mg/mL, GIBCO). Compounds were diluted to a final concentration of 0.5 μM and incubated with microsomal protein at 0.25 mg/mL in phosphate-buffered saline (PBS) at pH 7.4. The DMSO content in the incubation mixture was maintained below 1%. The metabolic reaction was initiated by introducing NADPH as a cofactor at a concentration of 0.5 μM. Aliquots were taken at defined time intervals: 0 (immediately after NADPH addition), 5, 10, 20, 30, and 60 min. Reactions were halted by the addition of a quenching solvent consisting of a 1:1 mixture of acetonitrile and methanol containing an internal standard at 50 nM. Following quenching, samples were centrifuged at 3500 rpm for 30 min to pellet the precipitated microsomal proteins. The resulting supernatants were analyzed via LC–MS/MS. Quantification was performed based on the peak area ratio (PAR) of the analyte to internal standard, with the signal at time zero defined as 100%. The percentage of parent compound remaining at each time point was calculated accordingly. Using the plot of % remaining versus incubation time, the degradation rate constant (k) was determined via nonlinear regression. From this, the half-life (*T*_1/2_ = ln(2)/k, in minutes) and intrinsic clearance (CL_int_ = k × 1000/0.25, in μL/min/mg protein) were calculated. Chromatographic analysis was carried out using a Supelco Ascentis Express C18 column (3 cm × 2.1 mm, 5 μm particle size). The mobile phases were composed of water with 0.1% formic acid (A) and acetonitrile with 0.1% formic acid (B). A binary gradient was applied as follows: 0–0.05 min, 95% A; 0.3:0.7 min, 2% A; 0.8:2.0 min, re-equilibration at 95% A. The total run time was 2 min per sample, with an injection volume of 10 μL and a flow rate of 0.7 mL/min. All metabolic stability assays were conducted in triplicate.

## Data availability

All datasets and source code used in this study are publicly available through GitHub. The repository for holistic GNN development and benchmarking is accessible at https://github.com/LCi-UFG/HolisticGNN. The repository containing the scripts and data for target fishing validation is available at https://github.com/LCi-UFG/BayesDocking.

## CRediT authorship contribution statement

**Vinícius Alexandre Fiaia Costa:** Data curation, Formal Analysis, Investigation, Software, Validation, Writing – original draft and Writing – review and editing. **Alexandra Maria dos Santos Carvalho:** Formal Analysis, Investigation, Validation, Visualization and Writing – review and editing. **Rafael Consolin Chelucci:** Formal Analysis, Investigation, Validation, Visualization and Writing- review and editing. **Felipe da Silva Mendonça de Melo:** Formal Analysis, Investigation and Validation. **Gustavo Santos Sandes Felizardo:** Formal Analysis, Investigation, Software, Visualization and Writing- review and editing. **Clarissa Alves Carneiro:** Data curation, Formal Analysis, Visualization. **Holli-Joi Martin:** Investigation, Validation, Writing – original draft and Writing – review and editing. **Rodolpho de Campos Braga:** Investigation, Supervision, validation and Visualization. **Sébastien Charneau:** Project administration, Resources and Supervision. **Eugene N. Muratov:** Supervision, Validation and Writing – review and editing. **Adriano Defini Andricopulo:** Methodology, Supervision and Validation. **Izabela Marques Dourado Bastos:** Conceptualization, Funding acquisition, Methodology, Project administration, Resources, Supervision, Visualization and Writing – review and editing. **Bruno Junior Neves:** Conceptualization, Funding acquisition, Investigation Methodology, Project administration, Resources, Software, Supervision, Validation, Writing – original draft and Writing – review and editing.

## Notes

Rodolpho de Campos Braga and Eugene N. Muratov are co-founders of InsilicAll LLC and Predictive LLC, respectively, which develop novel alternative methods and software for toxicity prediction. All the other authors declare no conflicts.

## REFERENCES

(1) Burza, S.; Croft, S. L.; Boelaert, M. Leishmaniasis. The Lancet 2018, 6736 (18), 1–20. 10.1016/S0140-6736(18)31204-2.

(2) Scarpini, S.; Dondi, A.; Totaro, C.; Biagi, C.; Melchionda, F.; Zama, D.; Pierantoni, L.; Gennari, M.; Campagna, C.; Prete, A.; Lanari, M. Visceral Leishmaniasis: Epidemiology, Diagnosis, and Treatment Regimens in Different Geographical Areas with a Focus on Pediatrics. Microorganisms 2022, 10 (10), 1887. 10.3390/microorganisms10101887.

(3) van Griensven, J.; Diro, E. Visceral Leishmaniasis. Infectious Disease Clinics of North America 2012, 26 (2), 309–322. 10.1016/j.idc.2012.03.005.

(4) Mann, S.; Frasca, K.; Scherrer, S.; Henao-Martínez, A. F.; Newman, S.; Ramanan, P.; Suarez, J. A. A Review of Leishmaniasis: Current Knowledge and Future Directions. Current Tropical Medicine Reports 2021, 8 (2), 121–132. 10.1007/s40475-021-00232-7.

(5) Word Health Organization. Leishmaniasis, 2025. https://www.who.int/news-room/fact-sheets/detail/leishmaniasis (accessed 2025-06-30).

(6) Dorlo, T. P. C.; Ostyn, B. A.; Uranw, S.; Dujardin, J.-C.; Boelaert, M. Treatment of Visceral Leishmaniasis: Pitfalls and Stewardship. The Lancet Infectious Diseases 2016, 16 (7), 777–778. 10.1016/S1473-3099(16)30091-3.

(7) Pradhan, S.; Schwartz, R. A.; Patil, A.; Grabbe, S.; Goldust, M. Treatment Options for Leishmaniasis. Clinical and Experimental Dermatology 2022, 47 (3), 516–521. 10.1111/ced.14919.

(8) Romero, G. A. S.; Costa, D. L.; Costa, C. H. N.; de Almeida, R. P.; de Melo, E. V.; de Carvalho, S. F. G.; Rabello, A.; de Carvalho, A. L.; Sousa, A. de Q.; Leite, R. D.; Lima, S. S.; Amaral, T. A.; Alves, F. P.; Rode, J. Efficacy and Safety of Available Treatments for Visceral Leishmaniasis in Brazil: A Multicenter, Randomized, Open Label Trial. PLOS Neglected Tropical Diseases 2017, 11 (6), e0005706. 10.1371/journal.pntd.0005706.

(9) Yang, X.; Wang, Y.; Byrne, R.; Schneider, G.; Yang, S. Concepts of Artificial Intelligence for Computer-Assisted Drug Discovery. Chemical Reviews 2019, 119 (18), 10520–10594. 10.1021/acs.chemrev.8b00728.

(10) Ekins, S.; Puhl, A. C.; Zorn, K. M.; Lane, T. R.; Russo, D. P.; Klein, J. J.; Hickey, A. J.; Clark, A. M. Exploiting Machine Learning for End-to-End Drug Discovery and Development. Nature Materials 2019, 18 (5), 435–441. 10.1038/s41563-019-0338-z.

(11) Santos, E. S. de A.; Lemos, J. M.; dos Santos Carvalho, A. M.; Mendonça de Melo, F. da S.; Pereira, E. de S.; Moreira-Filho, J. T.; Braga, R. de C.; Muratov, E. N.; Grellier, P.; Charneau, S.; Bastos, I. M. D.; Neves, B. J. Deep Multitask Learning-Driven Discovery of New Compounds Targeting Leishmania Infantum. ACS Omega 2024, 9 (52), 51271–51284. 10.1021/acsomega.4c07994.

(12) Linciano, P.; Quotadamo, A.; Luciani, R.; Santucci, M.; Zorn, K. M.; Foil, D. H.; Lane, T. R.; Cordeiro da Silva, A.; Santarem, N.; B Moraes, C.; Freitas-Junior, L.; Wittig, U.; Mueller, W.; Tonelli, M.; Ferrari, S.; Venturelli, A.; Gul, S.; Kuzikov, M.; Ellinger, B.; Reinshagen, J.; Ekins, S.; Costi, M. P. High-Throughput Phenotypic Screening and Machine Learning Methods Enabled the Selection of Broad-Spectrum Low-Toxicity Antitrypanosomatidic Agents. J. Med. Chem. 2023, 66 (22), 15230–15255. 10.1021/acs.jmedchem.3c01322.

(13) Barros de Menezes, R. P.; Ferreira de Sousa, N.; Herrera-Acevedo, C.; Santos Aquino de Araújo, R.; Durand Trigueiro Lira, N.; Fechine Tavares, J.; Jorge Kato, M.; Alex da Rocha Coelho, F.; Sousa dos Santos, A. L.; Antonio da Franca Rodrigues, K.; Bezerra Mendonça-Júnior, F. J.; Scotti, L.; Tullius Scotti, M. MuDRA-Based Virtual Screening of Terpenes for Anti-Leishmania Infantum Activity: In Vitro Validation and Mechanistic Insights from Molecular Docking. ChemMedChem 2025, 20 (4), e202400743. 10.1002/cmdc.202400743.

(14) Grisoni, F.; Consonni, V.; Todeschini, R. Impact of Molecular Descriptors on Computational Models. In Computational Chemogenomics; Humana Press: New York, 2018; Vol. 1825, pp 171–209. 10.1007/978-1-4939-8639-2_5.

(15) Rogers, D.; Hahn, M. Extended-Connectivity Fingerprints. J. Chem. Inf. Model. 2010, 50 (5), 742–754. 10.1021/ci100050t.

(16) Zhang, O.; Lin, H.; Zhang, X.; Wang, X.; Wu, Z.; Ye, Q.; Zhao, W.; Wang, J.; Ying, K.; Kang, Y.; Hsieh, C.; Hou, T. Graph Neural Networks in Modern AI-Aided Drug Discovery. arXiv June 7, 2025. 10.48550/arXiv.2506.06915.

(17) Wieder, O.; Kohlbacher, S.; Kuenemann, M.; Garon, A.; Ducrot, P.; Seidel, T.; Langer, T. A Compact Review of Molecular Property Prediction with Graph Neural Networks. Drug Discovery Today: Technologies 2020, 37, 1–12. 10.1016/j.ddtec.2020.11.009.

(18) Zhou, J.; Cui, G.; Hu, S.; Zhang, Z.; Yang, C.; Liu, Z.; Wang, L.; Li, C.; Sun, M. Graph Neural Networks: A Review of Methods and Applications. AI Open 2020, 1, 57–81. 10.1016/j.aiopen.2021.01.001.

(19) Khemani, B.; Patil, S.; Kotecha, K.; Tanwar, S. A Review of Graph Neural Networks: Concepts, Architectures, Techniques, Challenges, Datasets, Applications, and Future Directions. Journal of Big Data 2024, 11 (1), 18. 10.1186/s40537-023-00876-4.

(20) Deng, J.; Yang, Z.; Wang, H.; Ojima, I.; Samaras, D.; Wang, F. A Systematic Study of Key Elements Underlying Molecular Property Prediction. Nat Commun 2023, 14 (1), 6395. 10.1038/s41467-023-41948-6.

(21) Gilmer, J.; Schoenholz, S. S.; Riley, P. F.; Vinyals, O.; Dahl, G. E. Neural Message Passing for Quantum Chemistry. arXiv June 12, 2017. 10.48550/arXiv.1704.01212.

(22) Veličković, P.; Cucurull, G.; Casanova, A.; Romero, A.; Liò, P.; Bengio, Y. Graph Attention Networks. arXiv February 4, 2018. 10.48550/arXiv.1710.10903.

(23) Xu, K.; Hu, W.; Leskovec, J.; Jegelka, S. How Powerful Are Graph Neural Networks? arXiv February 22, 2019. 10.48550/arXiv.1810.00826.

(24) Xiong, Z.; Wang, D.; Liu, X.; Zhong, F.; Wan, X.; Li, X.; Li, Z.; Luo, X.; Chen, K.; Jiang, H.; Zheng, M. Pushing the Boundaries of Molecular Representation for Drug Discovery with the Graph Attention Mechanism. J Med Chem 2020, 63 (16), 8749–8760. 10.1021/acs.jmedchem.9b00959.

(25) Liu, C.; Sun, Y.; Davis, R.; Cardona, S. T.; Hu, P. ABT-MPNN: An Atom-Bond Transformer-Based Message-Passing Neural Network for Molecular Property Prediction. Journal of Cheminformatics 2023, 15 (1), 29. 10.1186/s13321-023-00698-9.

(26) Ying, C.; Cai, T.; Luo, S.; Zheng, S.; Ke, G.; He, D.; Shen, Y.; Liu, T.-Y. Do Transformers Really Perform Bad for Graph Representation?

(27) Yang, K.; Swanson, K.; Jin, W.; Coley, C.; Eiden, P.; Gao, H.; Guzman-Perez, A.; Hopper, T.; Kelley, B.; Mathea, M.; Palmer, A.; Settels, V.; Jaakkola, T.; Jensen, K.; Barzilay, R. Analyzing Learned Molecular Representations for Property Prediction. J. Chem. Inf. Model. 2019, 59 (8), 3370–3388. 10.1021/acs.jcim.9b00237.

(28) Examining graph neural networks for crystal structures: Limitations and opportunities for capturing periodicity | Science Advances. https://www.science.org/doi/10.1126/sciadv.adi3245 (accessed 2025-06-26).

(29) Hierarchical Molecular Graph Self-Supervised Learning for property prediction | Communications Chemistry. https://www.nature.com/articles/s42004-023-00825-5 (accessed 2025-06-26).

(30) Banjade, H. R.; Hauri, S.; Zhang, S.; Ricci, F.; Gong, W.; Hautier, G.; Vucetic, S.; Yan, Q. Structure Motif–Centric Learning Framework for Inorganic Crystalline Systems. Science Advances 2021, 7 (17), eabf1754. 10.1126/sciadv.abf1754.

(31) Chen, J.; Schwaller, P. Molecular Hypergraph Neural Networks. The Journal of Chemical Physics 2024, 160 (14), 144307. 10.1063/5.0193557.

(32) Reiser, P.; Neubert, M.; Eberhard, A.; Torresi, L.; Zhou, C.; Shao, C.; Metni, H.; van Hoesel, C.; Schopmans, H.; Sommer, T.; Friederich, P. Graph Neural Networks for Materials Science and Chemistry. Commun Mater 2022, 3 (1), 93. 10.1038/s43246-022-00315-6.

(33) Zhao, Y.; Li, H.; Zhou, H.; Attar, H. R.; Pfaff, T.; Li, N. A Review of Graph Neural Network Applications in Mechanics-Related Domains. Artif Intell Rev 2024, 57 (11), 315. 10.1007/s10462-024-10931-y.

(34) Alon, U.; Yahav, E. On the Bottleneck of Graph Neural Networks and Its Practical Implications. arXiv March 9, 2021. 10.48550/arXiv.2006.05205.

(35) Akansha, S. Over-Squashing in Graph Neural Networks: A Comprehensive Survey. Neurocomputing 2025, 642, 130389. 10.1016/j.neucom.2025.130389.

(36) Giraldo, J. H.; Skianis, K.; Bouwmans, T.; Malliaros, F. D. On the Trade-off between Over-Smoothing and Over-Squashing in Deep Graph Neural Networks. In Proceedings of the 32nd ACM International Conference on Information and Knowledge Management; 2023; pp 566–576. 10.1145/3583780.3614997.

(37) Gaulton, A.; Bellis, L. J.; Bento, A. P.; Chambers, J.; Davies, M.; Hersey, A.; Light, Y.; McGlinchey, S.; Michalovich, D.; Al-Lazikani, B.; Overington, J. P. ChEMBL: A Large-Scale Bioactivity Database for Drug Discovery. Nucleic Acids Research 2012, 40 (Database issue), D1100–D1107. 10.1093/nar/gkr777.

(38) Katsuno, K.; Burrows, J. N.; Duncan, K.; van Huijsduijnen, R. H.; Kaneko, T.; Kita, K.; Mowbray, C. E.; Schmatz, D.; Warner, P.; Slingsby, B. T. Hit and Lead Criteria in Drug Discovery for Infectious Diseases of the Developing World. Nature Reviews Drug Discovery 2015, 14 (11), 751–758. 10.1038/nrd4683.

(39) Chen, M.; Wei, Z.; Huang, Z.; Ding, B.; Li, Y. Simple and Deep Graph Convolutional Networks. arXiv July 4, 2020. 10.48550/arXiv.2007.02133.

(40) Li, Q.; Han, Z.; Wu, X. Deeper Insights Into Graph Convolutional Networks for Semi-Supervised Learning. Proceedings of the AAAI Conference on Artificial Intelligence 2018, 32 (1). 10.1609/aaai.v32i1.11604.

(41) Ying, C.; Cai, T.; Luo, S.; Zheng, S.; Ke, G.; He, D.; Shen, Y.; Liu, T.-Y. Do Transformers Really Perform Badly for Graph Representation? In Advances in Neural Information Processing Systems; Curran Associates, Inc., 2021; Vol. 34, pp 28877–28888.

(42) Xu, K.; Li, C.; Tian, Y.; Sonobe, T.; Kawarabayashi, K.; Jegelka, S. Representation Learning on Graphs with Jumping Knowledge Networks. arXiv June 25, 2018. 10.48550/arXiv.1806.03536.

(43) Käppel, M.; Ackermann, L.; Jablonski, S.; Härtl, S. Explaining Transformer-Based next Activity Prediction by Using Attention Scores. Process Sci 2025, 2 (1), 11. 10.1007/s44311-025-00018-4.

(44) Serrano, S.; Smith, N. A. Is Attention Interpretable? arXiv June 9, 2019. 10.48550/arXiv.1906.03731.

(45) Jain, S.; Wallace, B. C. Attention Is Not Explanation. arXiv May 8, 2019. 10.48550/arXiv.1902.10186.

(46) P. Wellawatte, G.; Seshadri, A.; D. White, A. Model Agnostic Generation of Counterfactual Explanations for Molecules. Chemical Science 2022, 13 (13), 3697–3705. 10.1039/D1SC05259D.

(47) Thompson, A. M.; O’Connor, P. D.; Blaser, A.; Yardley, V.; Maes, L.; Gupta, S.; Launay, D.; Martin, D.; Franzblau, S. G.; Wan, B.; Wang, Y.; Ma, Z.; Denny, W. A. Repositioning Antitubercular 6-Nitro-2,3-Dihydroimidazo[2,1- *b*][1,3]Oxazoles for Neglected Tropical Diseases: Structure–Activity Studies on a Preclinical Candidate for Visceral Leishmaniasis. J. Med. Chem. 2016, 59 (6), 2530–2550. 10.1021/acs.jmedchem.5b01699.

(48) Thompson, A. M.; O’Connor, P. D.; Marshall, A. J.; Yardley, V.; Maes, L.; Gupta, S.; Launay, D.; Braillard, S.; Chatelain, E.; Franzblau, S. G.; Wan, B.; Wang, Y.; Ma, Z.; Cooper, C. B.; Denny, W. A. 7-Substituted 2-Nitro-5,6-Dihydroimidazo[2,1-b][1,3]Oxazines: Novel Antitubercular Agents Lead to a New Preclinical Candidate for Visceral Leishmaniasis. J Med Chem 2017, 60 (10), 4212–4233. 10.1021/acs.jmedchem.7b00034.

(49) Thompson, A. M.; O’Connor, P. D.; Marshall, A. J.; Blaser, A.; Yardley, V.; Maes, L.; Gupta, S.; Launay, D.; Braillard, S.; Chatelain, E.; Wan, B.; Franzblau, S. G.; Ma, Z.; Cooper, C. B.; Denny, W. A. Development of (6R)-2-Nitro-6-[4-(Trifluoromethoxy)Phenoxy]-6,7-Dihydro-5H-Imidazo[2,1-b][1,3]Oxazine (DNDI-8219): A New Lead for Visceral Leishmaniasis. J Med Chem 2018, 61 (6), 2329–2352. 10.1021/acs.jmedchem.7b01581.

(50) Thompson, A. M.; O’Connor, P. D.; Yardley, V.; Maes, L.; Launay, D.; Braillard, S.; Chatelain, E.; Wan, B.; Franzblau, S. G.; Ma, Z.; Cooper, C. B.; Denny, W. A. Novel Linker Variants of Antileishmanial/Antitubercular 7-Substituted 2-Nitroimidazooxazines Offer Enhanced Solubility. ACS Med Chem Lett 2021, 12 (2), 275–281. 10.1021/acsmedchemlett.0c00649.

(51) Mowbray, C. E.; Braillard, S.; Glossop, P. A.; Whitlock, G. A.; Jacobs, R. T.; Speake, J.; Pandi, B.; Nare, B.; Maes, L.; Yardley, V.; Freund, Y.; Wall, R. J.; Carvalho, S.; Bello, D.; Van den Kerkhof, M.; Caljon, G.; Gilbert, I. H.; Corpas-Lopez, V.; Lukac, I.; Patterson, S.; Zuccotto, F.; Wyllie, S. DNDI-6148: A Novel Benzoxaborole Preclinical Candidate for the Treatment of Visceral Leishmaniasis. J. Med. Chem. 2021, 64 (21), 16159–16176. 10.1021/acs.jmedchem.1c01437.

(52) Emami, S.; Tavangar, P.; Keighobadi, M. An Overview of Azoles Targeting Sterol 14α-Demethylase for Antileishmanial Therapy. European Journal of Medicinal Chemistry 2017, 135, 241–259. 10.1016/j.ejmech.2017.04.044.

(53) Ashok, P.; Chander, S.; Smith, T. K.; Prakash Singh, R.; Jha, P. N.; Sankaranarayanan, M. Biological Evaluation and Structure Activity Relationship of 9-Methyl-1-Phenyl-*9H*-Pyrido[3,4-*b*]Indole Derivatives as Anti-Leishmanial Agents. Bioorganic Chemistry 2019, 84, 98–105. 10.1016/j.bioorg.2018.11.037.

(54) Upadhayaya, R. S.; Dixit, S. S.; Földesi, A.; Chattopadhyaya, J. New Antiprotozoal Agents: Their Synthesis and Biological Evaluations. Bioorganic & Medicinal Chemistry Letters 2013, 23 (9), 2750–2758. 10.1016/j.bmcl.2013.02.054.

(55) Di Giorgio, C.; Shimi, K.; Boyer, G.; Delmas, F.; Galy, J.-P. Synthesis and Antileishmanial Activity of 6-Mono-Substituted and 3,6-Di-Substituted Acridines Obtained by Acylation of Proflavine. Eur J Med Chem 2007, 42 (10), 1277–1284. 10.1016/j.ejmech.2007.02.010.

(56) Venkatraj, M.; Salado, I. G.; Heeres, J.; Joossens, J.; Lewi, P. J.; Caljon, G.; Maes, L.; Van der Veken, P.; Augustyns, K. Novel Triazine Dimers with Potent Antitrypanosomal Activity. European Journal of Medicinal Chemistry 2018, 143, 306–319. 10.1016/j.ejmech.2017.11.075.

(57) Berg, M.; Veken, P. V. der; Joossens, J.; Muthusamy, V.; Breugelmans, M.; Moss, C. X.; Rudolf, J.; Cos, P.; Coombs, G. H.; Maes, L.; Haemers, A.; Mottram, J. C.; Augustyns, K. Design and Evaluation of *Trypanosoma Brucei* Metacaspase Inhibitors. Bioorganic & Medicinal Chemistry Letters 2010, 20 (6), 2001–2006. 10.1016/j.bmcl.2010.01.099.

(58) Computer-aided discovery of two novel chalcone-like compounds active and selective against Leishmania infantum - ScienceDirect. https://www.sciencedirect.com/science/article/pii/S0960894X17303748?via%3Dihub (accessed 2025-10-16).

(59) Koovits, P. J.; Dessoy, M. A.; Matheeussen, A.; Maes, L.; Caljon, G.; Mowbray, C. E.; Kratz, J. M.; Dias, L. C. Structure-Activity Relationship of 4-Azaindole-2-Piperidine Derivatives as Agents against *Trypanosoma Cruzi*. Bioorganic & Medicinal Chemistry Letters 2020, 30 (1), 126779. 10.1016/j.bmcl.2019.126779.

(60) Exertier, C.; Salerno, A.; Antonelli, L.; Fiorillo, A.; Ocello, R.; Seghetti, F.; Caciolla, J.; Uliassi, E.; Masetti, M.; Fiorentino, E.; Orsini, S.; Di Muccio, T.; Ilari, A.; Bolognesi, M. L. Fragment Merging, Growing, and Linking Identify New Trypanothione Reductase Inhibitors for Leishmaniasis. J. Med. Chem. 2024, 67 (1), 402–419. 10.1021/acs.jmedchem.3c01439.

(61) Intakhan, N.; Saeung, A.; Rodrigues Oliveira, S. M.; Pereira, M. de L.; Chanmol, W. Synergistic Effects of Artesunate in Combination with Amphotericin B and Miltefosine against Leishmania Infantum: Potential for Dose Reduction and Enhanced Therapeutic Strategies. Antibiotics 2024, 13 (9), 806. 10.3390/antibiotics13090806.

(62) Daina, A.; Michielin, O.; Zoete, V. SwissADME: A Free Web Tool to Evaluate Pharmacokinetics, Drug-Likeness and Medicinal Chemistry Friendliness of Small Molecules. Scientific Reports 2017, 7 (1), 42717. 10.1038/srep42717.

(63) La Rosa, C.; Sharma, P.; Junaid Dar, M.; Jin, Y.; Qin, L.; Roy, A.; Kendall, A.; Wu, M.; Lin, Z.; Uchenik, D.; Li, J.; Basu, S.; Moitra, S.; Zhang, K.; Zhuo Wang, M.; Werbovetz, K. A. N-Substituted-4-(Pyridin-4-Ylalkyl)Piperazine-1-Carboxamides and Related Compounds as Leishmania CYP51 and CYP5122A1 Inhibitors. Bioorg Med Chem 2024, 113, 117907. 10.1016/j.bmc.2024.117907.

(64) Jin, Y.; Basu, S.; Feng, M.; Ning, Y.; Munasinghe, I.; Joachim, A. M.; Li, J.; Qin, L.; Madden, R.; Burks, H.; Gao, P.; Wu, J. Q.; Sheikh, S. W.; Joice, A. C.; Perera, C.; Werbovetz, K. A.; Zhang, K.; Wang, M. Z. CYP5122A1 Encodes an Essential Sterol C4-Methyl Oxidase in Leishmania Donovani and Determines the Antileishmanial Activity of Antifungal Azoles. Nat Commun 2024, 15, 9409. 10.1038/s41467-024-53790-5.

(65) Hargrove, T. Y.; Kim, K.; de Nazaré Correia Soeiro, M.; da Silva, C. F.; da Gama Jaen Batista, D.; Batista, M. M.; Yazlovitskaya, E. M.; Waterman, M. R.; Sulikowski, G. A.; Lepesheva, G. I. CYP51 Structures and Structure-Based Development of Novel, Pathogen-Specific Inhibitory Scaffolds. Int J Parasitol Drugs Drug Resist 2012, 2, 178–186. 10.1016/j.ijpddr.2012.06.001.

(66) Lepesheva, G. I.; Hargrove, T. Y.; Rachakonda, G.; Wawrzak, Z.; Pomel, S.; Cojean, S.; Nde, P. N.; Nes, W. D.; Locuson, C. W.; Calcutt, M. W.; Waterman, M. R.; Daniels, J. S.; Loiseau, P. M.; Villalta, F. VFV as a New Effective CYP51 Structure-Derived Drug Candidate for Chagas Disease and Visceral Leishmaniasis. J Infect Dis 2015, 212 (9), 1439–1448. 10.1093/infdis/jiv228.

(67) Hargrove, T. Y.; Wawrzak, Z.; Liu, J.; Nes, W. D.; Waterman, M. R.; Lepesheva, G. I. Substrate Preferences and Catalytic Parameters Determined by Structural Characteristics of Sterol 14α-Demethylase (CYP51) from Leishmania Infantum*. Journal of Biological Chemistry 2011, 286 (30), 26838–26848. 10.1074/jbc.M111.237099.

(68) CYP5122A1 encodes an essential sterol C4-methyl oxidase in Leishmania donovani and determines the antileishmanial activity of antifungal azoles *- PMC*. https://pmc.ncbi.nlm.nih.gov/articles/PMC11528044/ (accessed 2025-06-29).

(69) Fourches, D.; Muratov, E.; Tropsha, A. Trust, but Verify: On the Importance of Chemical Structure Curation in Cheminformatics and QSAR Modeling Research. Journal of Chemical Information and Modeling 2010, 50 (7), 1189–1204. 10.1021/ci100176x.

(70) Fourches, D.; Muratov, E.; Tropsha, A. Trust, but Verify II: A Practical Guide to Chemogenomics Data Curation. Journal of Chemical Information and Modeling 2016, 56 (7), 1243–1252. 10.1021/acs.jcim.6b00129.

(71) Moreira-Filho, J. T.; Braga, R. C.; Lemos, J. M.; Alves, V. M.; Borba, J. V. V. B.; Costa, W. S.; Kleinstreuer, N.; Muratov, E. N.; Andrade, C. H.; Neves, B. J. BeeToxAI: An Artificial Intelligence-Based Web App to Assess Acute Toxicity of Chemicals to Honey Bees. Artificial Intelligence in the Life Sciences 2021, 1, 100013. 10.1016/j.ailsci.2021.100013.

(72) Shelley, J. C.; Cholleti, A.; Frye, L. L.; Greenwood, J. R.; Timlin, M. R.; Uchimaya, M. Epik: A Software Program for pK (a) Prediction and Protonation State Generation for Drug-like Molecules. Journal of computer-aided molecular design 2007, 21 (12), 681–691. 10.1007/s10822-007-9133-z.

(73) Søndergaard, C. R.; Olsson, M. H. M.; Rostkowski, M.; Jensen, J. H. Improved Treatment of Ligands and Coupling Effects in Empirical Calculation and Rationalization of pKa Values. Journal of Chemical Theory and Computation 2011, 7 (7), 2284–2295. 10.1021/ct200133y.

(74) Olsson, M. H. M.; Søndergaard, C. R.; Rostkowski, M.; Jensen, J. H. PROPKA3: Consistent Treatment of Internal and Surface Residues in Empirical pKa Predictions. J Chem Theory Comput 2011, 7 (2), 525–537. 10.1021/ct100578z.

(75) Banks, J. L.; Beard, H. S.; Cao, Y.; Cho, A. E.; Damm, W.; Farid, R.; Felts, A. K.; Halgren, T. A.; Mainz, D. T.; Maple, J. R.; Murphy, R.; Philipp, D. M.; Repasky, M. P.; Zhang, L. Y.; Berne, B. J.; Friesner, R. A.; Gallicchio, E.; Levy, R. M. Integrated Modeling Program, Applied Chemical Theory (IMPACT). Journal of Computational Chemistry 2005, 26 (16), 1752–1780. 10.1002/jcc.20292.

(76) Halgren, T. A.; Murphy, R. B.; Friesner, R. A.; Beard, H. S.; Frye, L. L.; Pollard, W. T.; Banks, J. L. Glide: A New Approach for Rapid, Accurate Docking and Scoring. 2. Enrichment Factors in Database Screening. Journal of Medicinal Chemistry 2004, 47 (7), 1750–1759. 10.1021/jm030644s.

(77) Friesner, R. A.; Murphy, R. B.; Repasky, M. P.; Frye, L. L.; Greenwood, J. R.; Halgren, T. A.; Sanschagrin, P. C.; Mainz, D. T. Extra Precision Glide: Docking and Scoring Incorporating a Model of Hydrophobic Enclosure for Protein−Ligand Complexes. Journal of Medicinal Chemistry 2006, 49 (21), 6177–6196. 10.1021/jm051256o.

