## Supplementary information for "Semantic-Aware Graph Embedding Approach Uncovers LC-61, a Potent Anti–*Leishmania infantum* Compound"

### Support Information


**Table S1. Performance metrics for holistic GNN architectures on the unbalanced dataset.**

| Architecture | Set | Threshold | Accuracy | Recall | Specificity | G-mean | AUC |
|---|---|---|---|---|---|---|---|
| **MPNN** | Training | 0.500 | 0.8436 | 0.5775 | 0.8946 | 0.7188 | 0.8740 |
| | Validation | 0.500 | 0.8322 | 0.5417 | 0.8908 | 0.6946 | 0.8606 |
| | Test | 0.500 | 0.8542 | 0.5000 | 0.9113 | 0.6750 | 0.8589 |
| **GIN** | Training | 0.500 | 0.8780 | 0.3850 | 0.9724 | 0.6119 | 0.8853 |
| | Validation | 0.500 | 0.8671 | 0.3333 | 0.9748 | 0.5700 | 0.8400 |
| | Test | 0.500 | 0.9028 | 0.4000 | 0.9839 | 0.6273 | 0.9117 |
| **GAT** | Training | 0.500 | 0.8672 | 0.2798 | 0.9683 | 0.5205 | 0.8747 |
| | Validation | 0.500 | 0.8392 | 0.2500 | 0.9580 | 0.4894 | 0.8319 |
| | Test | 0.500 | 0.8611 | 0.2000 | 0.9677 | 0.4399 | 0.8927 |
| **AttentiveFP** | Training | 0.500 | 0.8368 | 0.5187 | 0.8976 | 0.6824 | 0.8165 |
| | Validation | 0.500 | 0.8182 | 0.5000 | 0.8824 | 0.6642 | 0.8141 |
| | Test | 0.500 | 0.8264 | 0.3500 | 0.9032 | 0.5623 | 0.7827 |

**Table S2. Performance metrics for calibrated holistic GNN architectures on the unbalanced dataset.**

| Architecture | Set | Threshold | Accuracy | Recall | Specificity | G-mean | AUC |
|---|---|---|---|---|---|---|---|
| **MPNN** | Training | 0.170 | 0.8058 | 0.8289 | 0.8014 | 0.8150 | 0.8740 |
| | Validation | 0.210 | 0.8182 | 0.8750 | 0.8067 | 0.8402 | 0.8606 |
| | Test | 0.210 | 0.7986 | 0.8500 | 0.7903 | 0.8196 | 0.8589 |
| **GIN** | Training | 0.130 | 0.7981 | 0.8503 | 0.7881 | 0.8186 | 0.8853 |
| | Validation | 0.110 | 0.7692 | 0.8333 | 0.7563 | 0.7939 | 0.8400 |
| | Test | 0.110 | 0.7708 | 0.9000 | 0.7500 | 0.8216 | 0.9117 |
| **GAT** | Training | 0.120 | 0.7921 | 0.8274 | 0.7861 | 0.8065 | 0.8747 |
| | Validation | 0.130 | 0.8042 | 0.8333 | 0.7983 | 0.8156 | 0.8319 |
| | Test | 0.130 | 0.8403 | 0.9000 | 0.8306 | 0.8646 | 0.8927 |
| **AttentiveFP** | Training | 0.250 | 0.7912 | 0.6738 | 0.8137 | 0.7405 | 0.8165 |
| | Validation | 0.270 | 0.7902 | 0.7083 | 0.8067 | 0.7559 | 0.8141 |
| | Test | 0.270 | 0.7847 | 0.6500 | 0.8065 | 0.7240 | 0.7827 |

**Table S3. Performance metrics for holistic GNN architectures on the balanced dataset.**

| Architecture | Set | Threshold | Accuracy | Recall | Specificity | G-mean | AUC |
|---|---|---|---|---|---|---|---|
| **MPNN** | Training | 0.500 | 0.8026 | 0.7655 | 0.8347 | 0.7993 | 0.8864 |
| | Validation | 0.500 | 0.7958 | 0.7931 | 0.7976 | 0.7954 | 0.8561 |
| | Test | 0.500 | 0.8156 | 0.7794 | 0.8493 | 0.8136 | 0.9158 |
| **GIN** | Training | 0.500 | 0.8330 | 0.7411 | 0.9125 | 0.8223 | 0.9306 |
| | Validation | 0.500 | 0.8169 | 0.7069 | 0.8929 | 0.7945 | 0.8734 |
| | Test | 0.500 | 0.8227 | 0.7647 | 0.8767 | 0.8188 | 0.9124 |
| **GAT** | Training | 0.500 | 0.8207 | 0.7301 | 0.8963 | 0.8089 | 0.9045 |
| | Validation | 0.500 | 0.8310 | 0.7759 | 0.8690 | 0.8211 | 0.8592 |
| | Test | 0.500 | 0.8511 | 0.7941 | 0.9041 | 0.8473 | 0.8902 |
| **AttentiveFP** | Training | 0.500 | 0.7939 | 0.6792 | 0.8930 | 0.7788 | 0.8782 |
| | Validation | 0.500 | 0.7746 | 0.6379 | 0.8690 | 0.7446 | 0.8177 |
| | Test | 0.500 | 0.7730 | 0.6618 | 0.8767 | 0.7617 | 0.8668 |

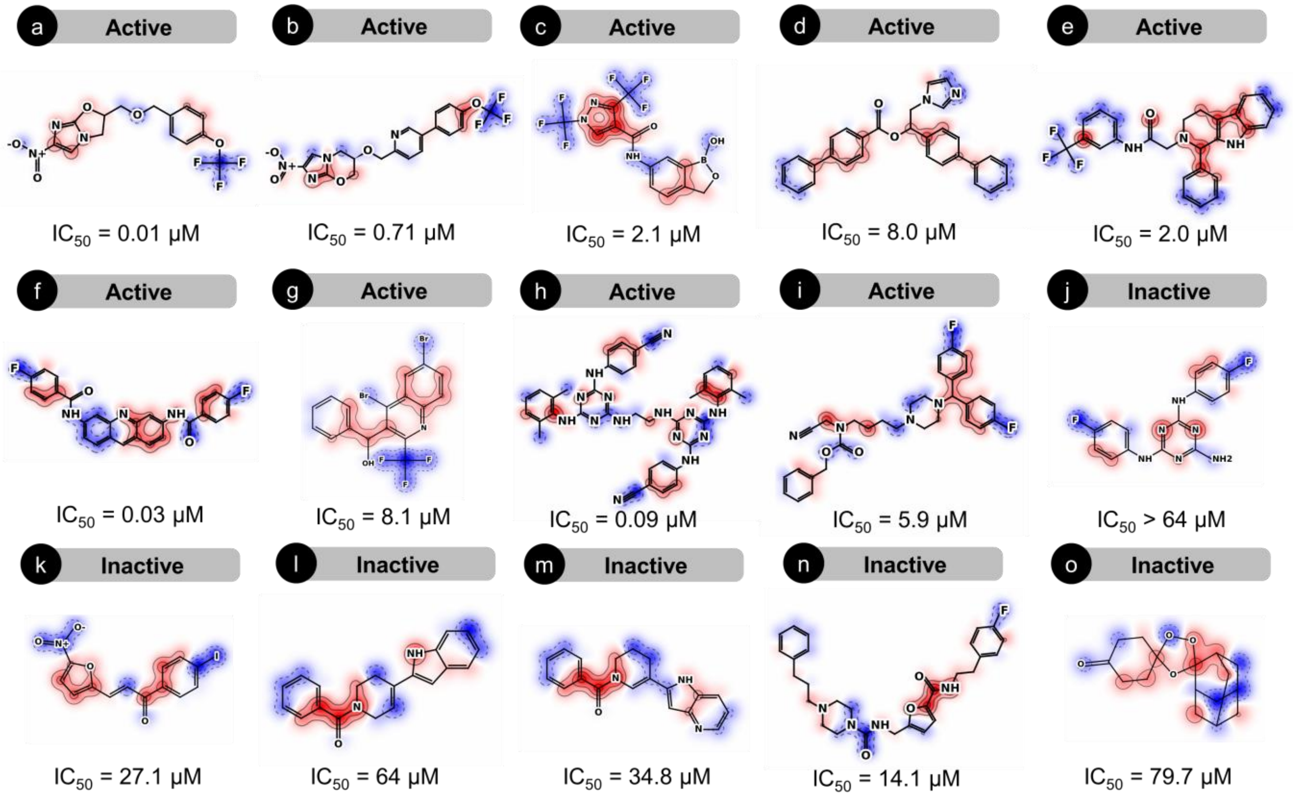


**Figure S1. Attentional explainability of predicted antileishmanial activity.** Red contours indicate regions contributing positively to antileishmanial activity, whereas blue contours indicate regions contributing negatively.

**LC-58** ({1-[(2-methylphenyl)methyl]piperidin-4-yl}methyl)({[5-(2-methylpropyl)-1H-pyrazol-3-yl]methyl})[(pyridin-3-yl)methyl]amine

ST290610

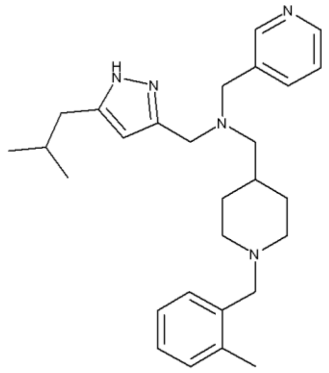

| ID | 91952146 | 445.6565 | $C_{28}H_{39}N_5$ |
|---|---|---|---|

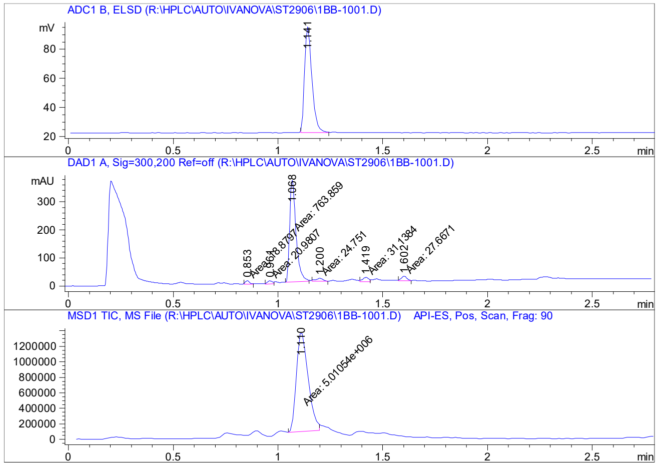


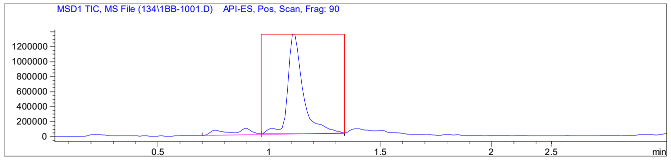


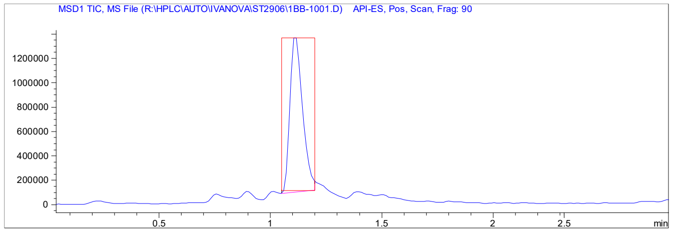


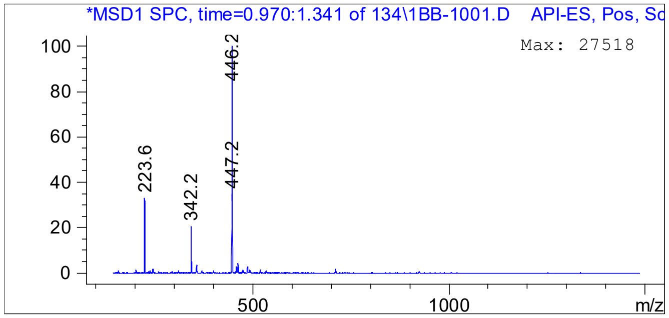


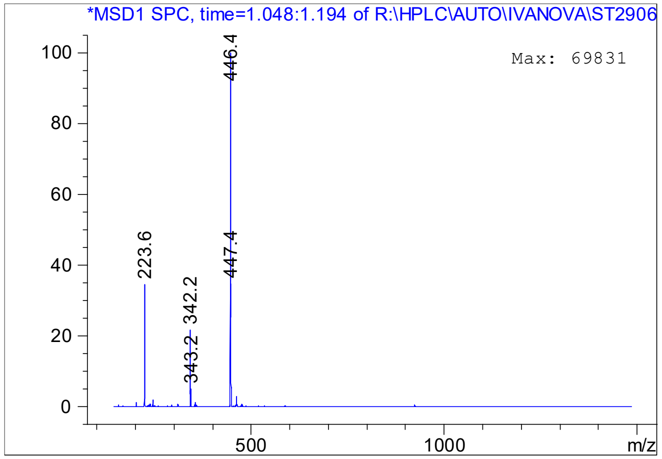

# LC-59

#### 2-ethyl-4-methyl-N-[2-(piperidin-1-yl)-2-(pyridin-3-yl)ethyl]-1,3-thiazole-5-carboxamide

ST836939

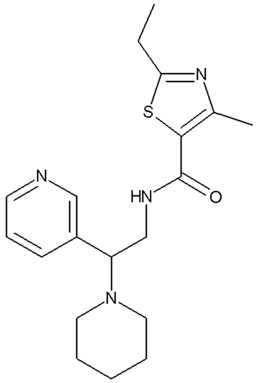

| | | | |
|---|---|---|---|
| ID | 78768815 | 358.5093 | $C_{19}H_{26}N_4OS$ |

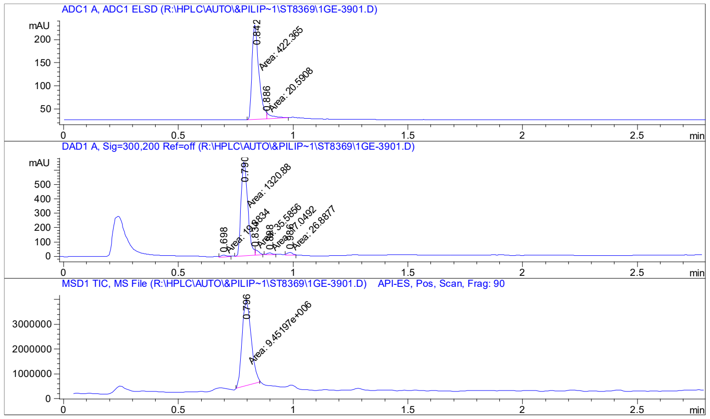


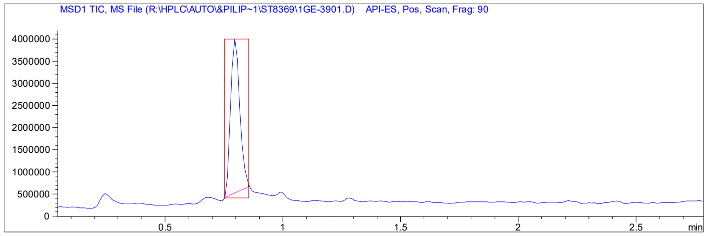


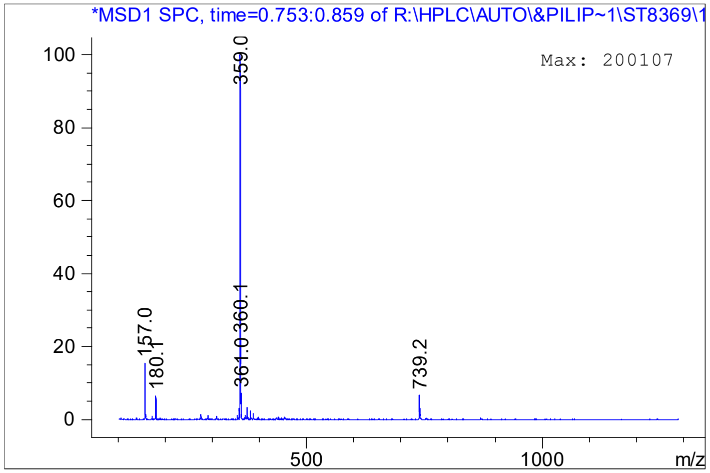

# LC-60

#### 1-(4-fluoro-2-methylphenyl)-N-[2-(1H-imidazol-1-yl)-1-phenylethyl]piperidin-4-amine

FC942392915

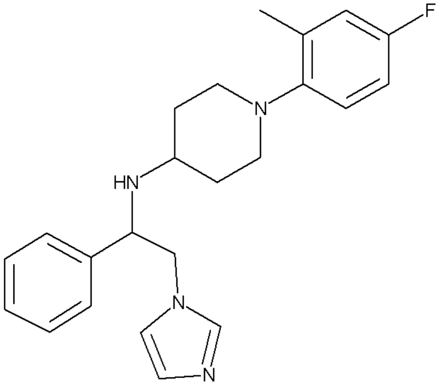

| | | | |
|---|---|---|---|
| ID | 73055423 | 378.4968 | $C_{23}H_{27}FN_4$ |

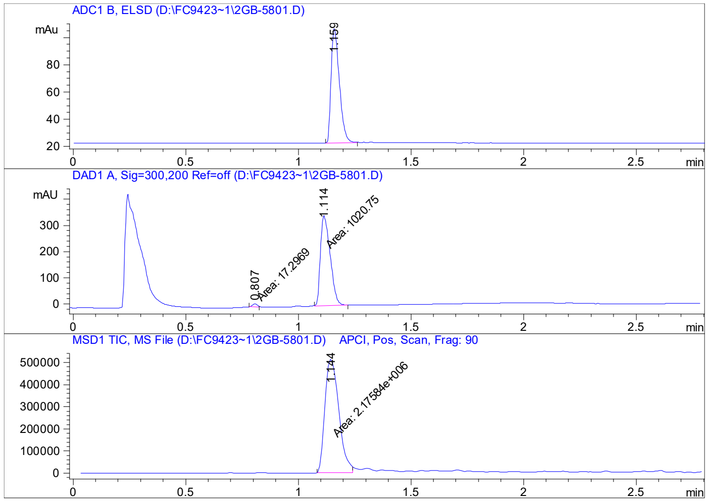


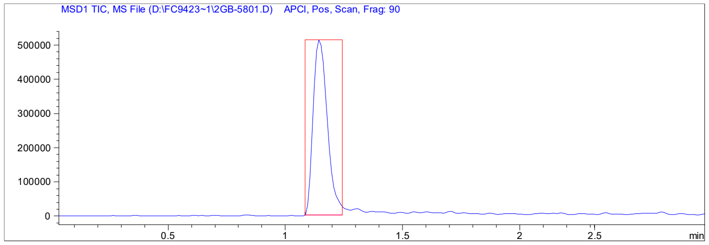


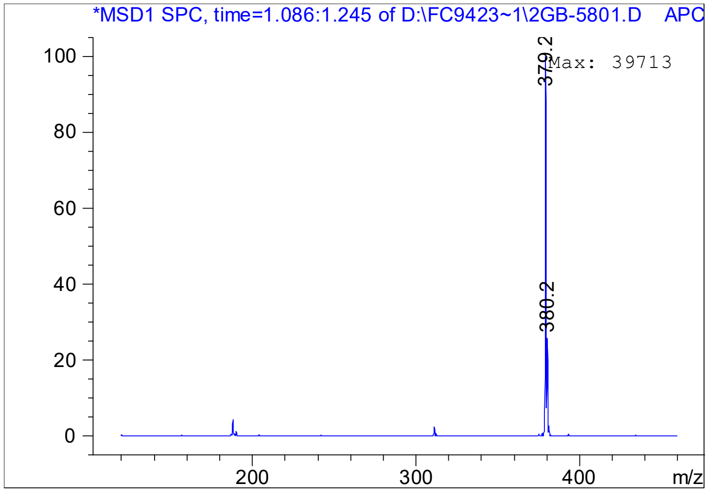

# LC-61

### 1-(2-fluorophenyl)-N-[1-(pyridin-3-yl)pentyl]piperidin-4-amine

FC942290706

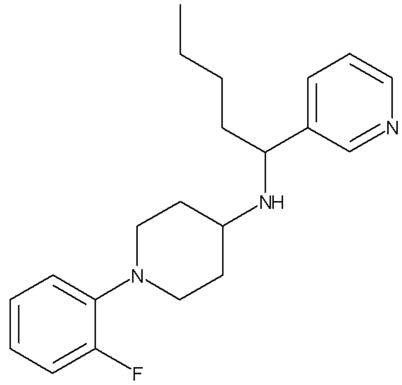


| | | | |
|---|---|---|---|
| ID | 49635071 | 341.4758 | $C_{21}H_{28}FN_3$ |

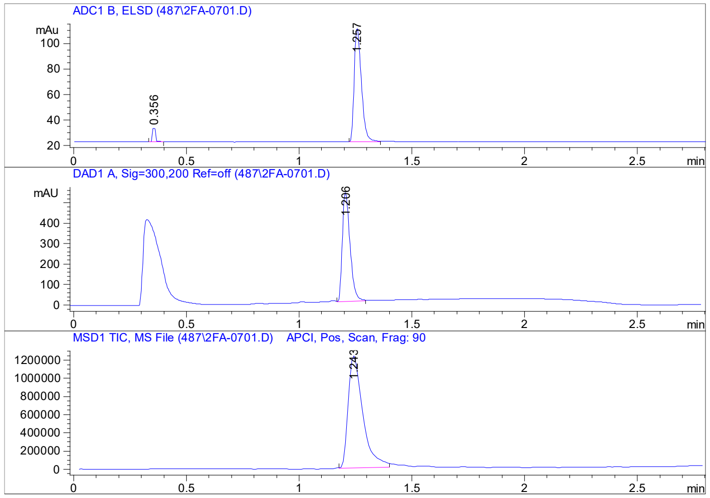


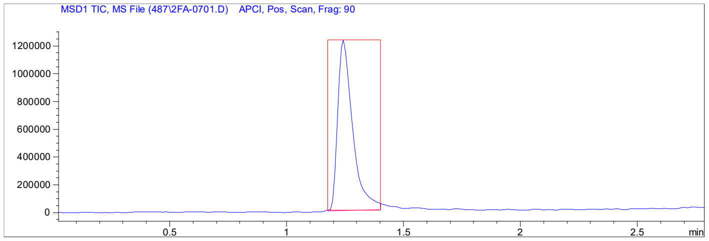


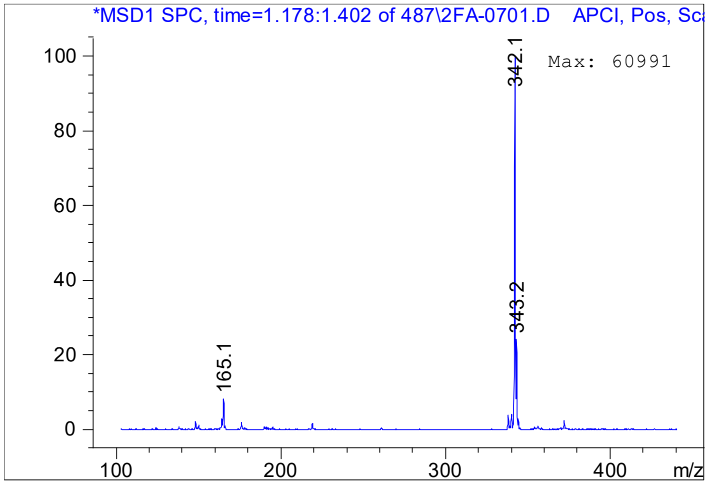

# LC-62

#### 1-(3,4-difluorophenyl)-N-[2-(piperidin-1-yl)-2-(pyridin-3-yl)ethyl]piperidin-4-amine

FC942433206

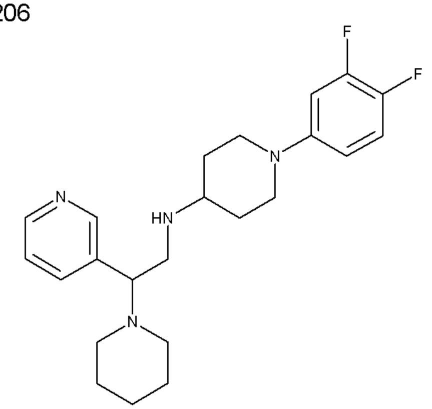


| | | | |
|---|---|---|---|
| ID | 46177884 | 400.5192 | $C_{23}H_{30}F_2N_4$ |

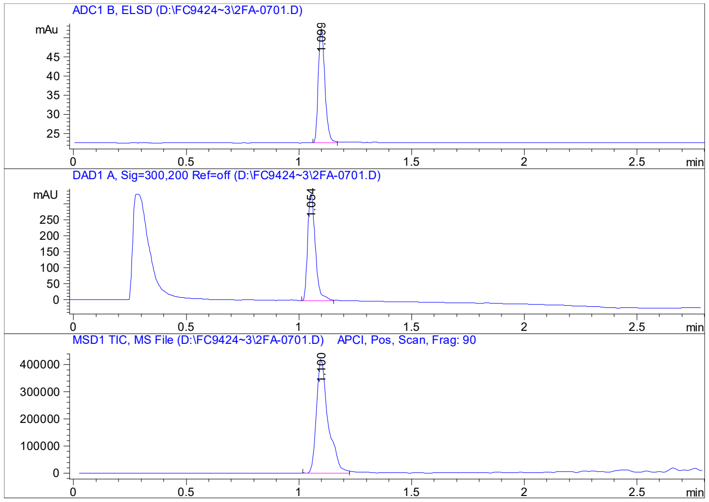


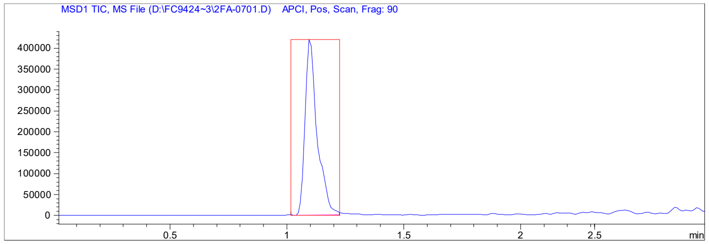


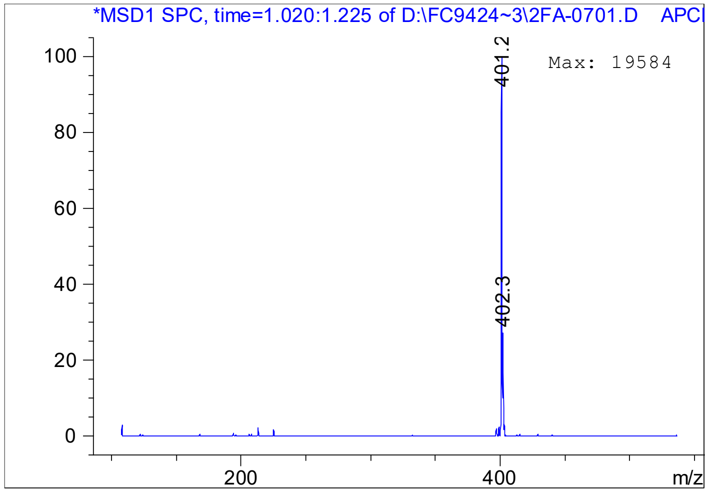

# LC-63

#### [(1-cyclohexylpiperidin-3-yl)methyl][(3-fluorophenyl)methyl][(pyridin-3-yl)methyl]amine

ST155310

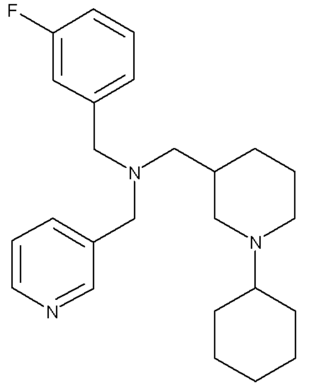

| | | | |
|---|---|---|---|
| ID | 33806354 | 395.5682 | $C_{25}H_{34}FN_3$ |

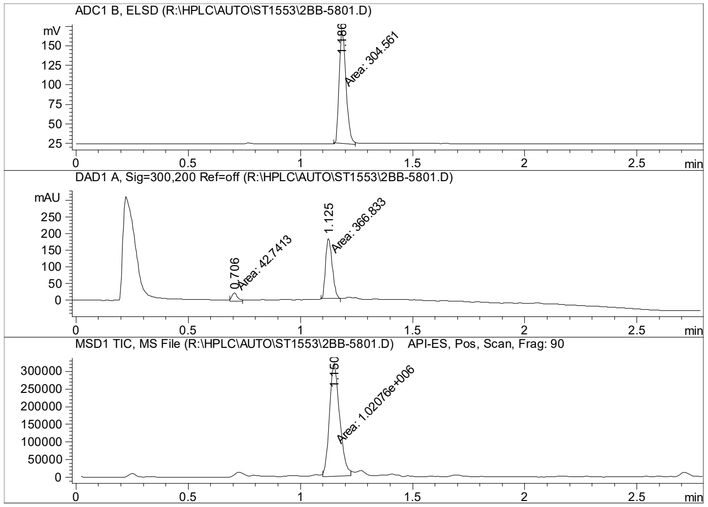


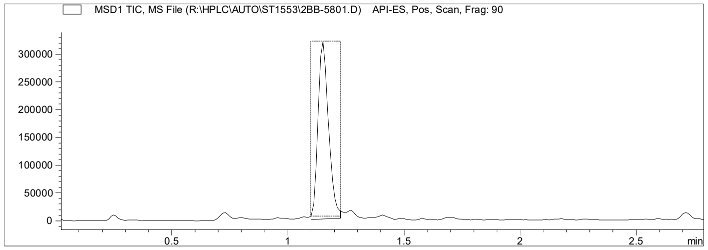


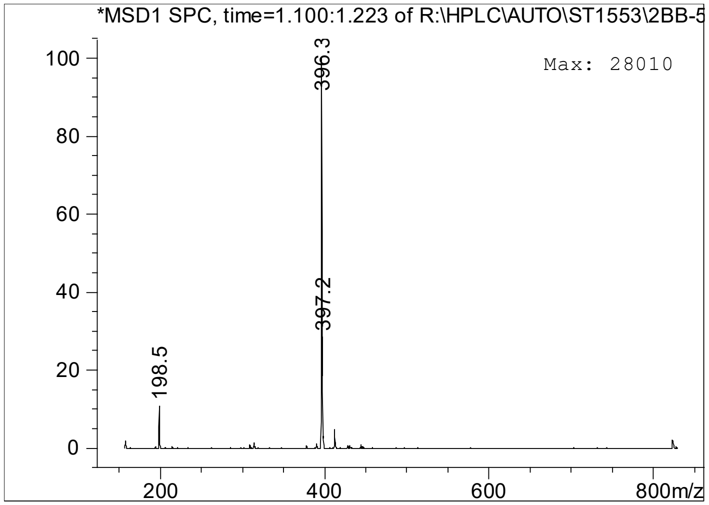

# LC-64

#### ({1-[(2-methylphenyl)methyl]piperidin-4-yl}methyl)({[4-(propan-2-yl)-1,3-thiazol-2-yl]methyl})[(pyridin-3-yl)methyl]amine

ST709407

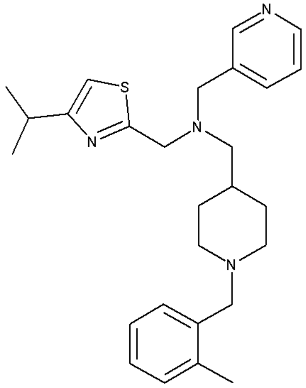


| | | | |
|---|---|---|---|
| ID | 27722619 | 448.6788 | $C_{27}H_{36}N_4S$ |

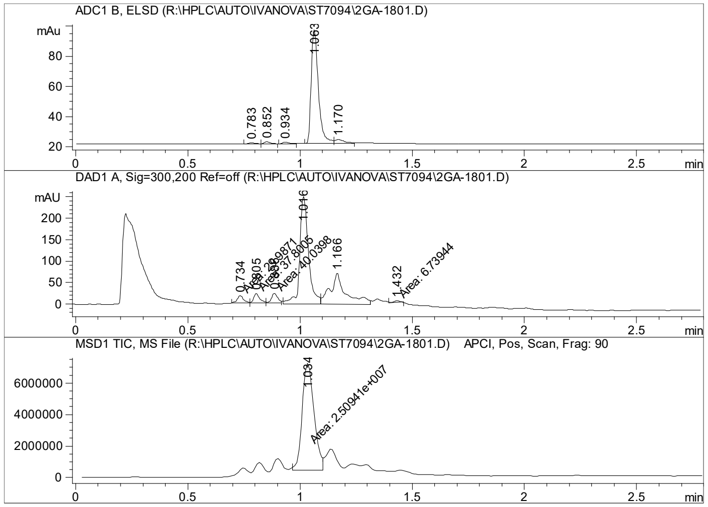


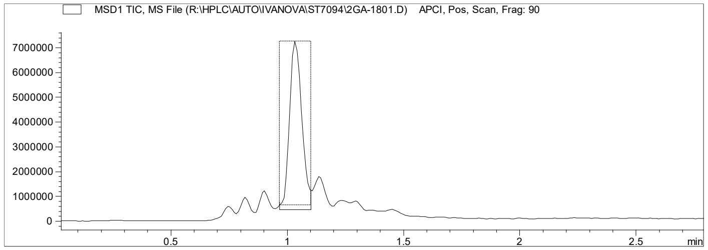


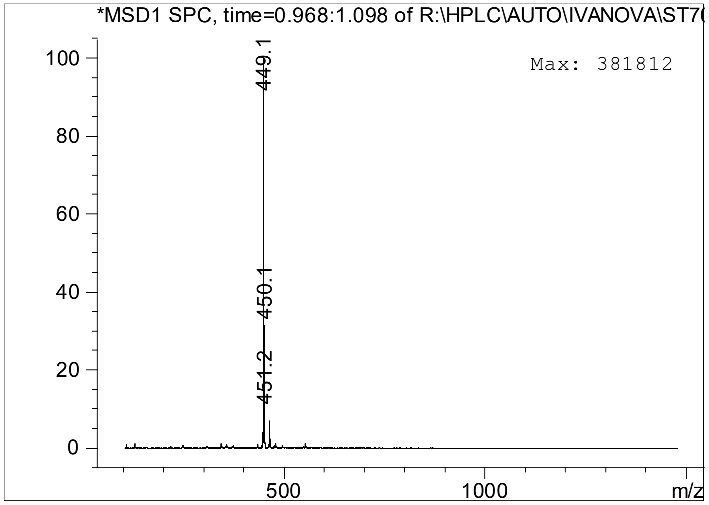

# LC-65

#### 1-{4-[(4-fluorophenyl)methyl]-1,4-diazepan-1-yl}-2-(morpholin-4-yl)-2-(pyridin-3-yl)ethan-1-one

FC942419920

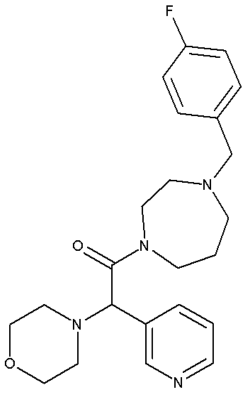

| | | | |
|---|---|---|---|
| ID | 25286552 | 412.5116 | $C_{23}H_{29}FN_4O_2$ |

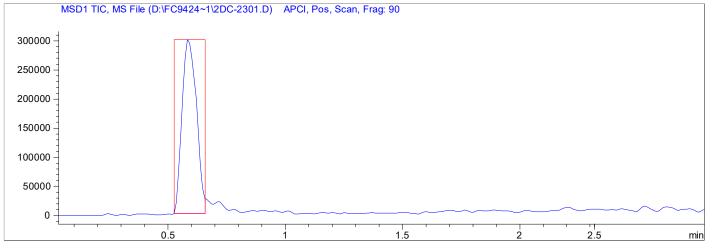


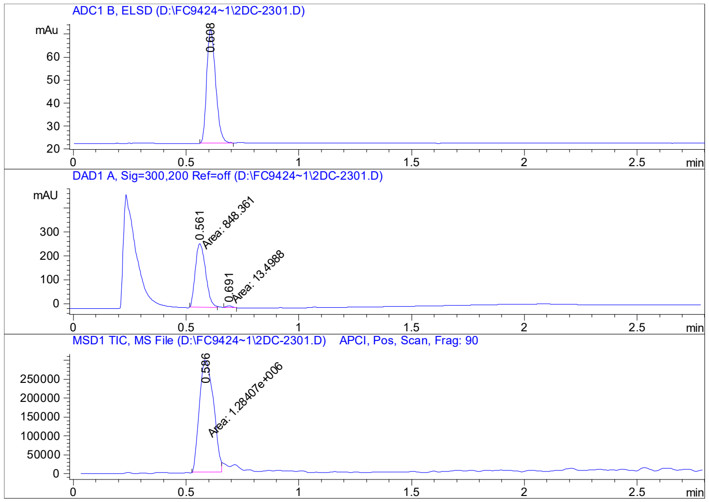


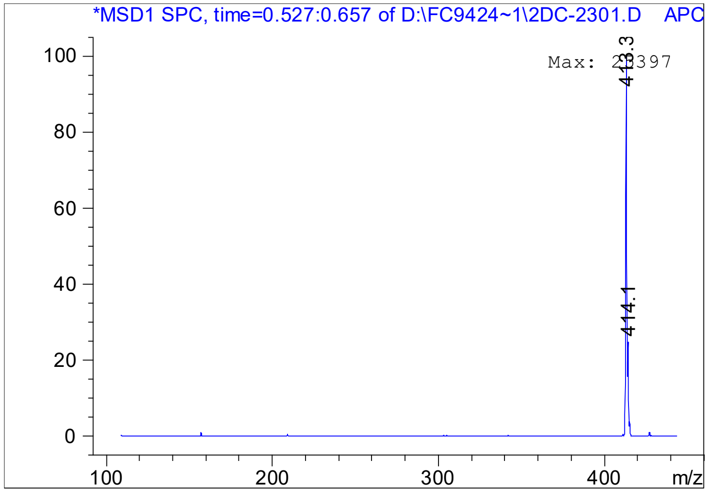

# LC-66

#### N-[(4-chlorophenyl)(pyridin-3-yl)methyl]-2-phenylacetamide

CH00704234

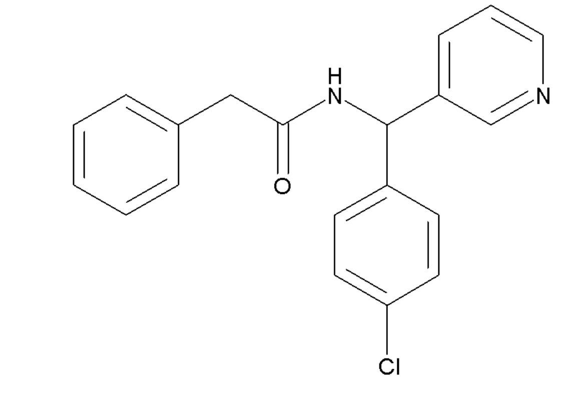


| | | | |
|---|---|---|---|
| ID | 9294189 | 336.8243 | $C_{20}H_{17}ClN_2O$ |

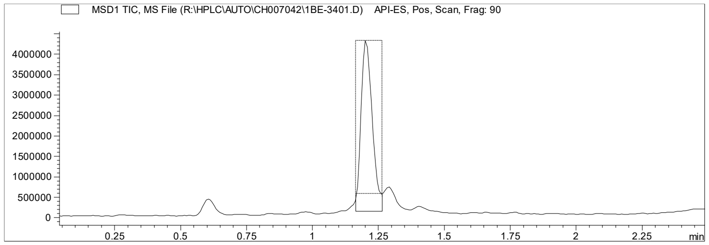


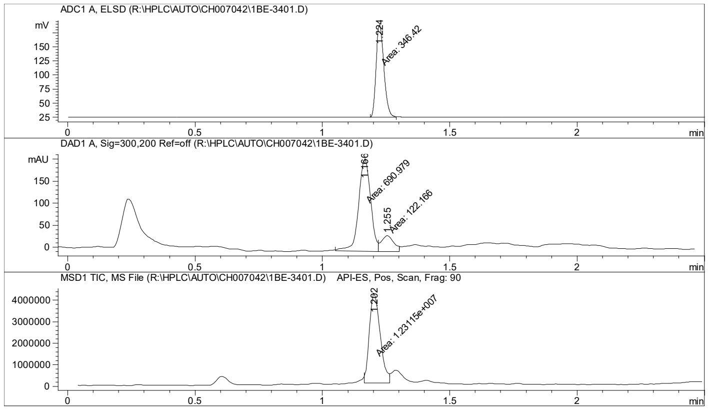


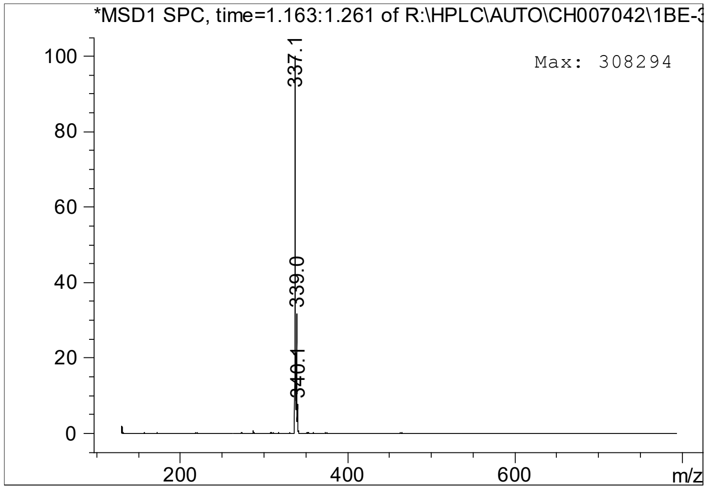

# LC-67

#### N-[2-(1H-imidazol-5-yl)ethyl]-2-phenylquinoline-4-carboxamide

CH00654570

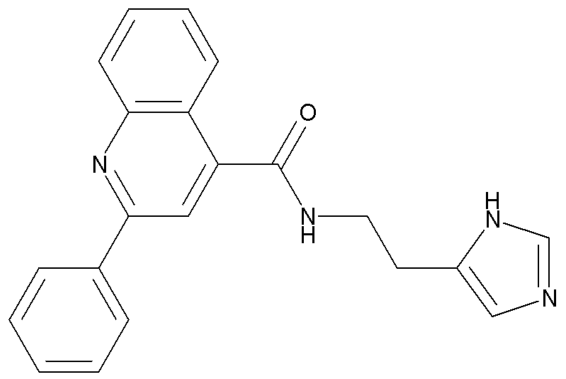

| | | | |
|---|---|---|---|
| ID | 9251367 | 342.4038 | $C_{21}H_{18}N_4O$ |

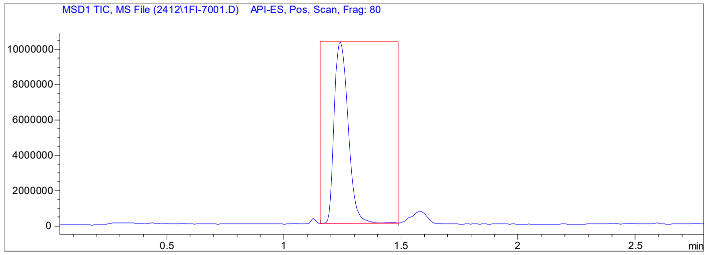


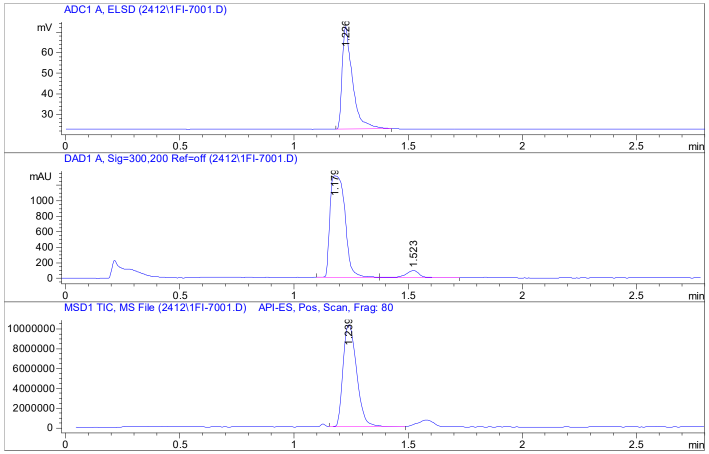


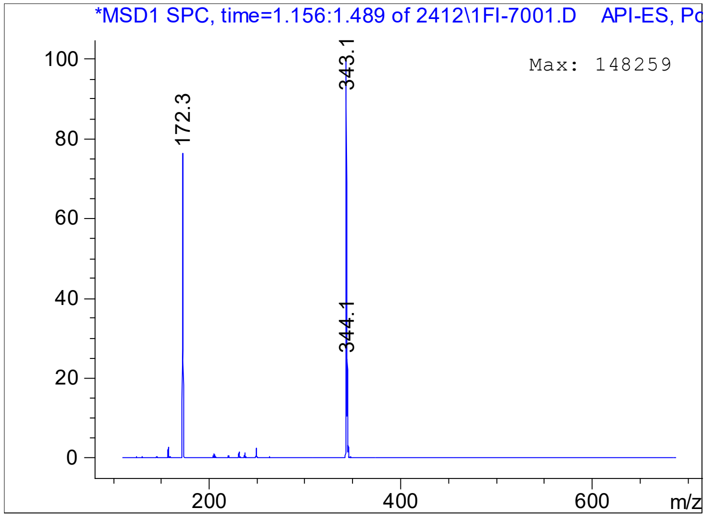

**LC-68**

**N-[4-(2,5-dimethylphenyl)-5-methyl-1,3-thiazol-2-yl]pyridine-4-carboxamide**

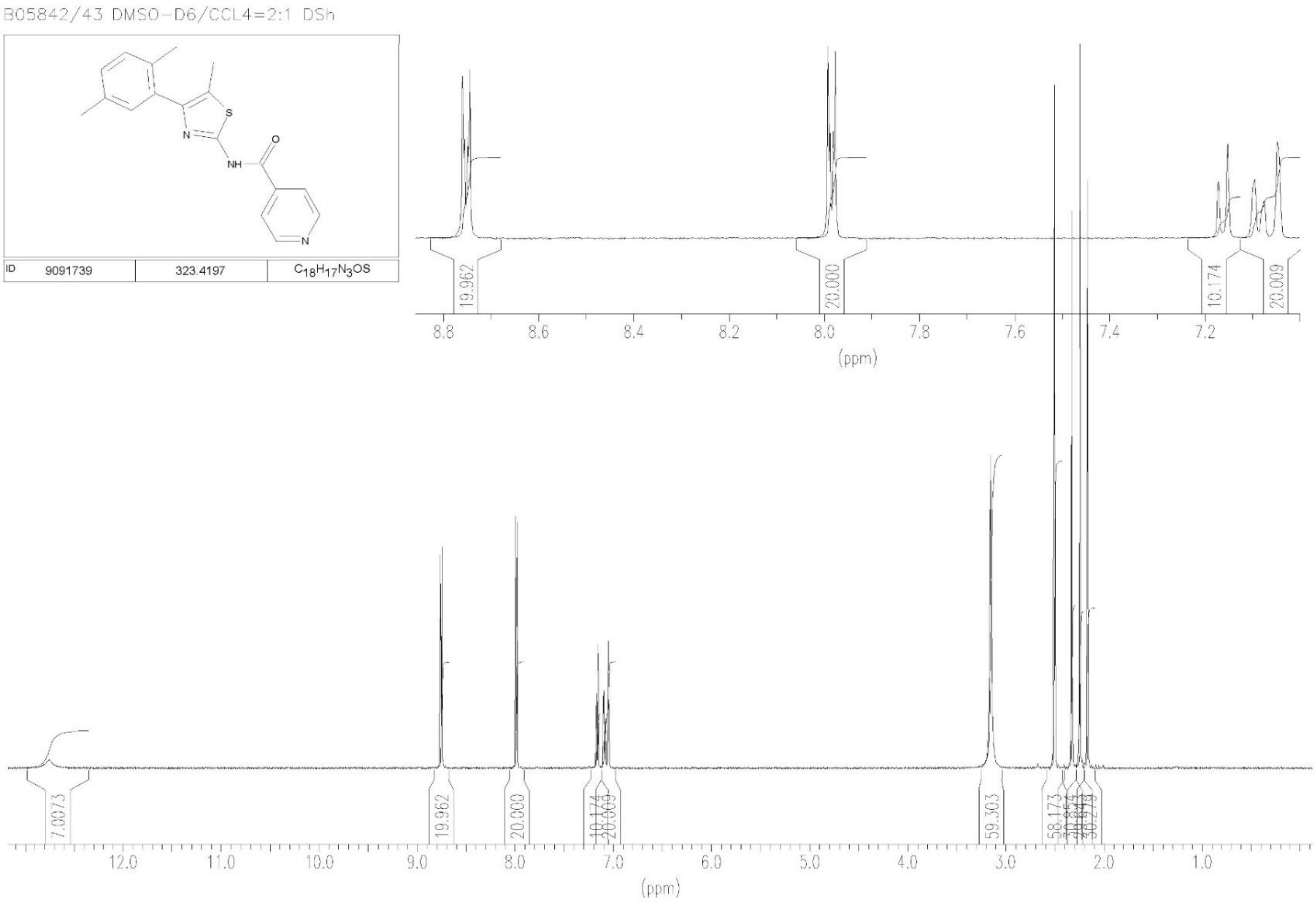

**LC-69**

**N-[4-(2,5-dimethylphenyl)-5-methyl-1,3-thiazol-2-yl]cyclopropanecarboxamide**

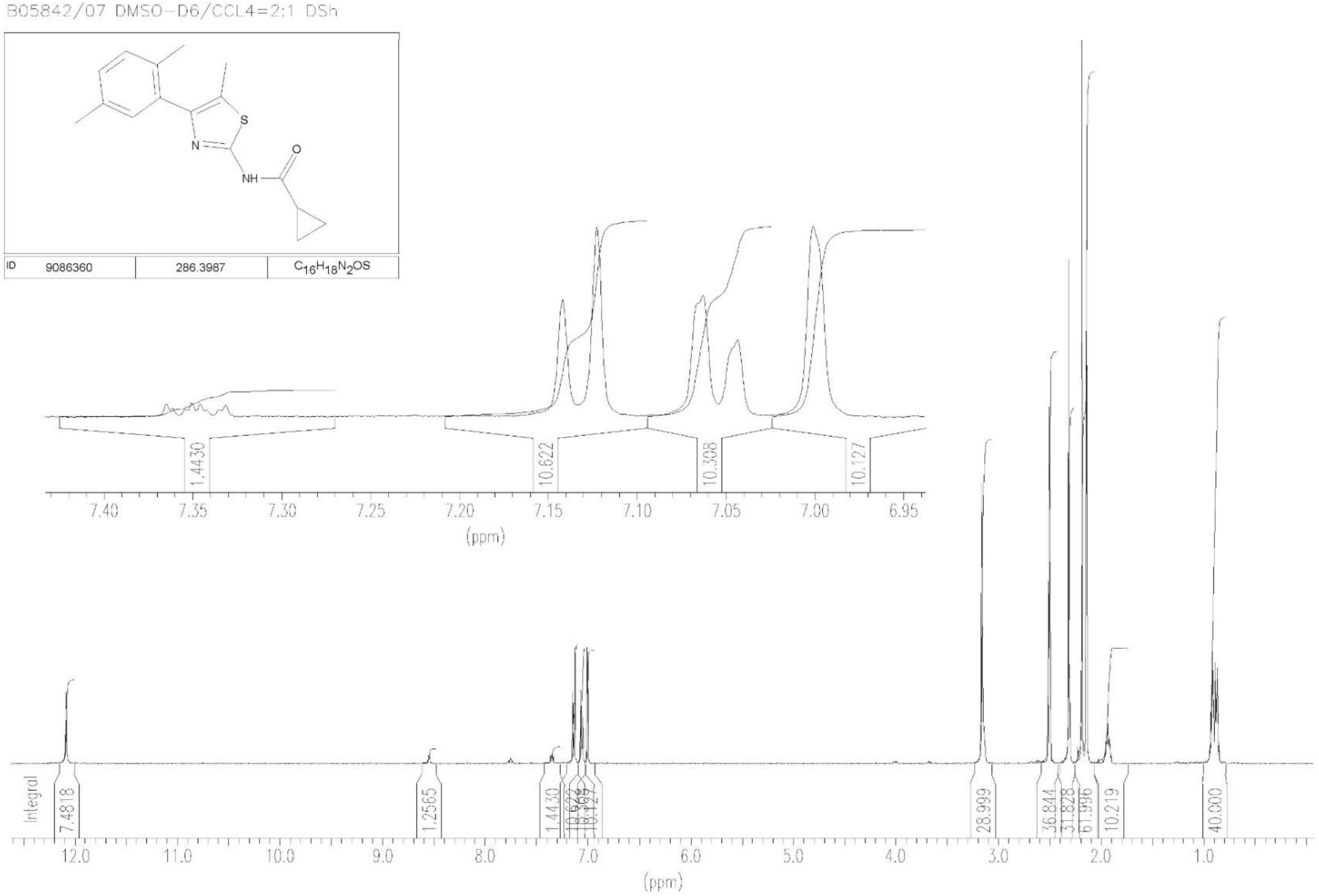

**LC-70**

**N-[(4-chlorophenyl)(pyridin-3-yl)methyl]-1-(propan-2-yl)piperidine-4-carboxamide**

FC941284303

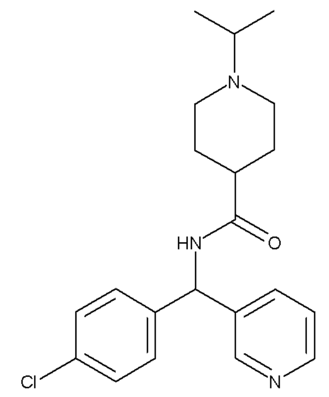

| | | | |
|---|---|---|---|
| ID | 39760868 | 371.9139 | $C_{21}H_{26}ClN_3O$ |

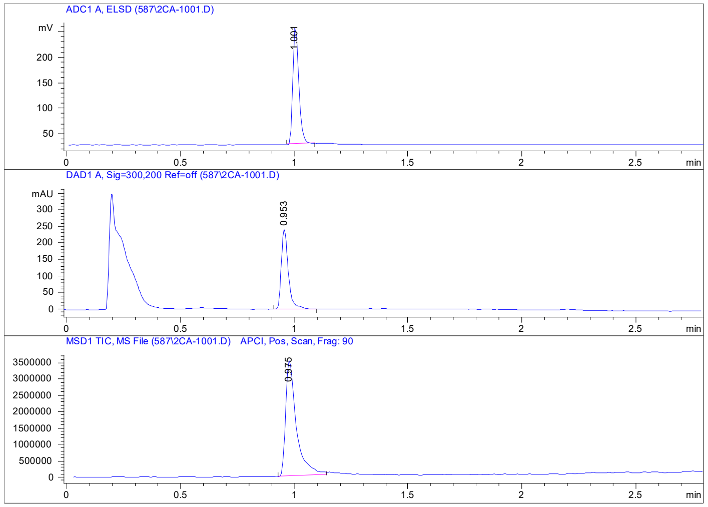


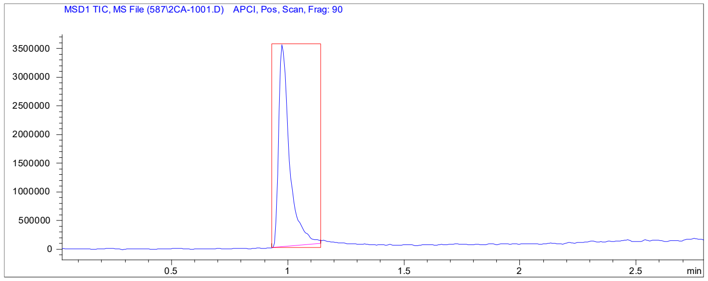


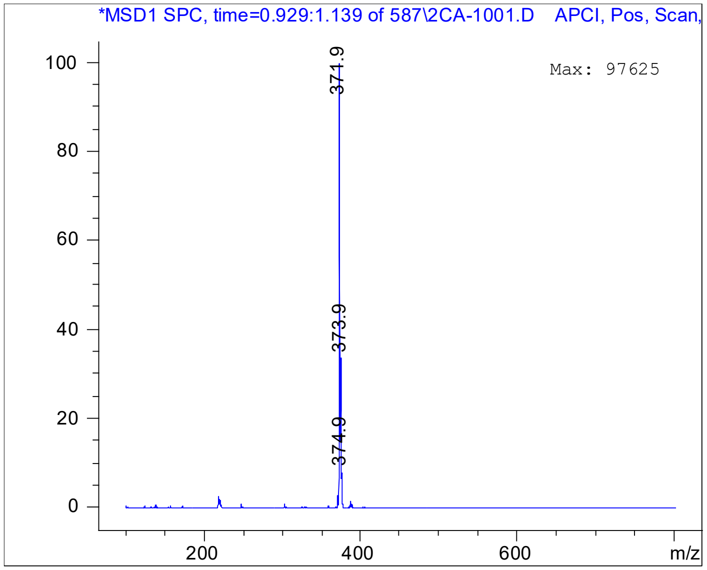

# LC-71

#### N-(3-oxo-1,3-dihydro-2-benzofuran-5-yl)-2-(pyridin-3-yl)piperidine-1-carboxamide

ST577528

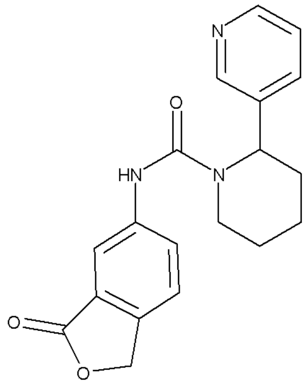

| | | | |
|---|---|---|---|
| ID | 59553984 | 337.3816 | $C_{19}H_{19}N_3O_3$ |

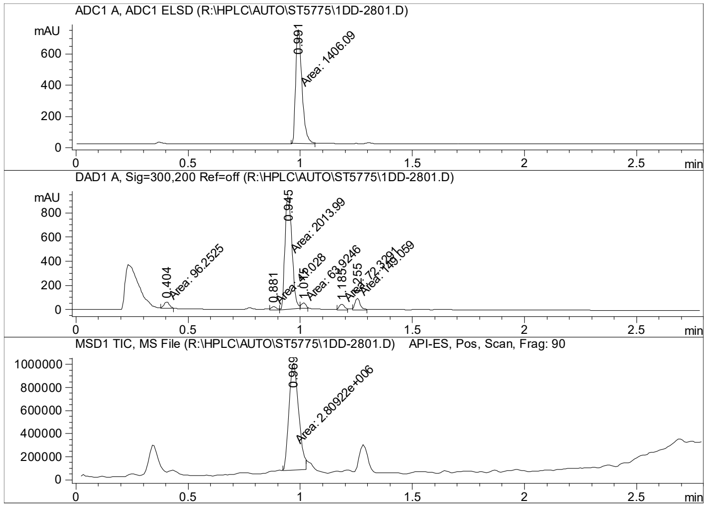


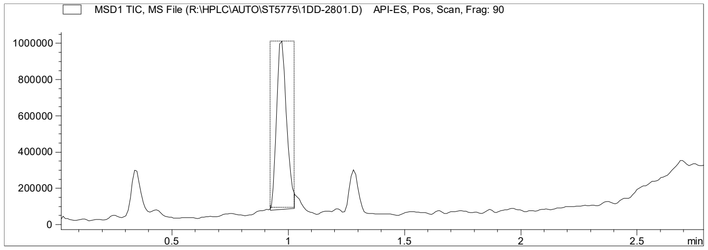


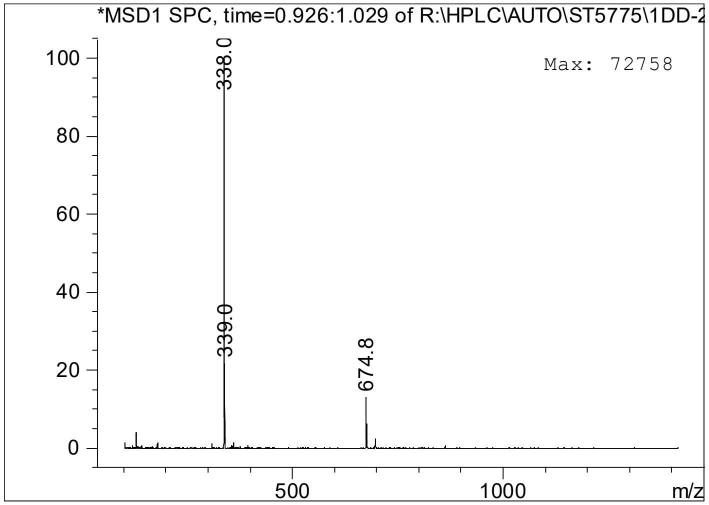

# LC-72

### N-(1-methyl-1H-1,2,3-benzotriazol-5-yl)-2-(pyridin-3-yl)piperidine-1-carboxamide

ST993926

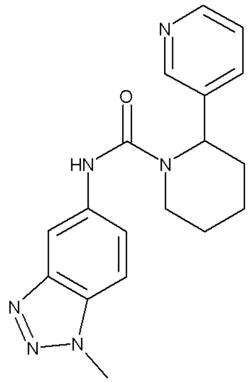


| | | | |
|---|---|---|---|
| ID | 74022226 | 336.3997 | $C_{18}H_{20}N_6O$ |

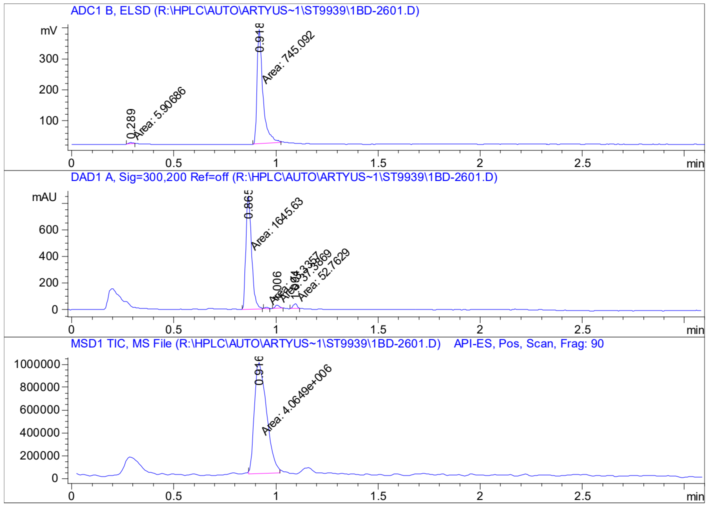


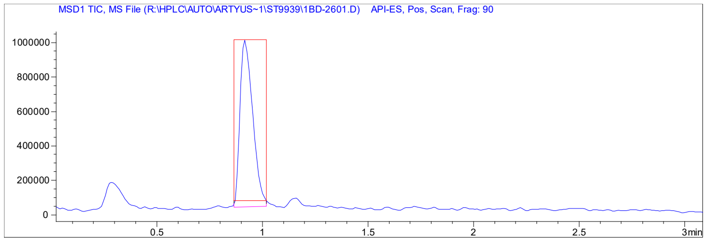


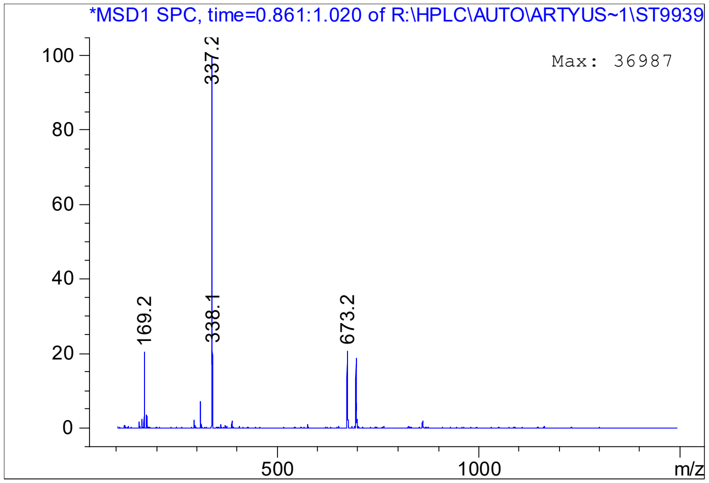

# LC-73

#### N-[(4-chlorophenyl)(pyridin-3-yl)methyl]-1-(propan-2-yl)-1H-pyrazole-4-carboxamide

ST892328

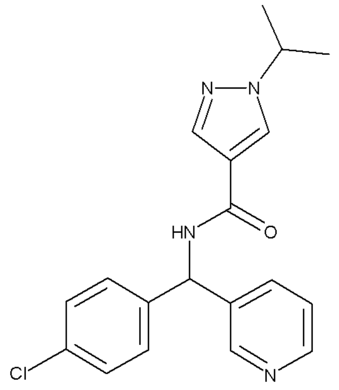

| ID | 13179027 | 354.8425 | $C_{19}H_{19}ClN_4O$ |
|---|---|---|---|

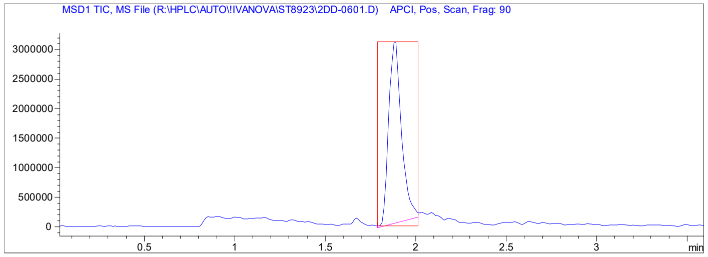


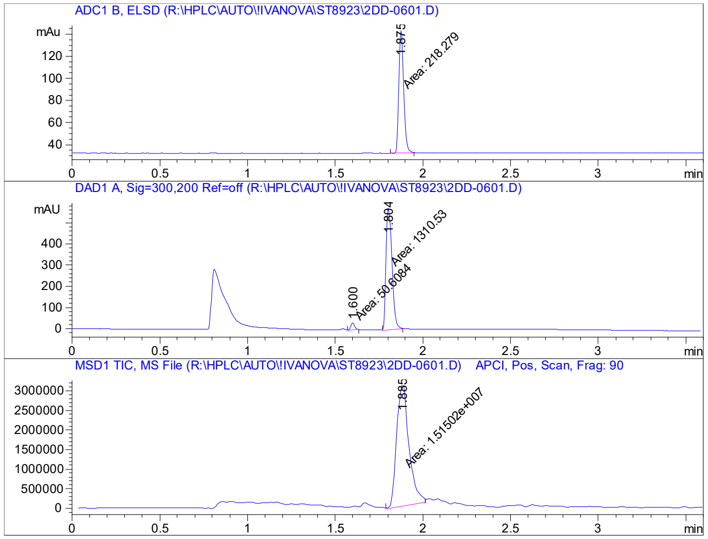


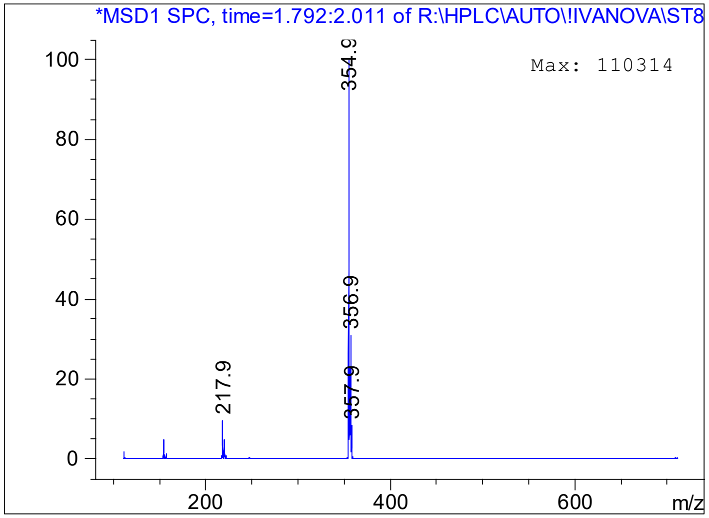

# LC-74

### N-(6-chloropyridin-3-yl)-2-(pyridin-3-yl)piperidine-1-carboxamide

ST642304

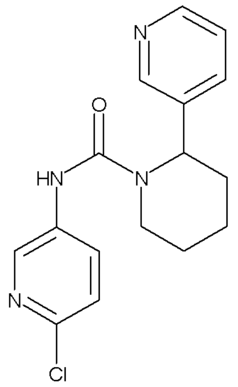


| | | | |
|---|---|---|---|
| ID | 79118598 | 316.7931 | $C_{16}H_{17}ClN_4O$ |

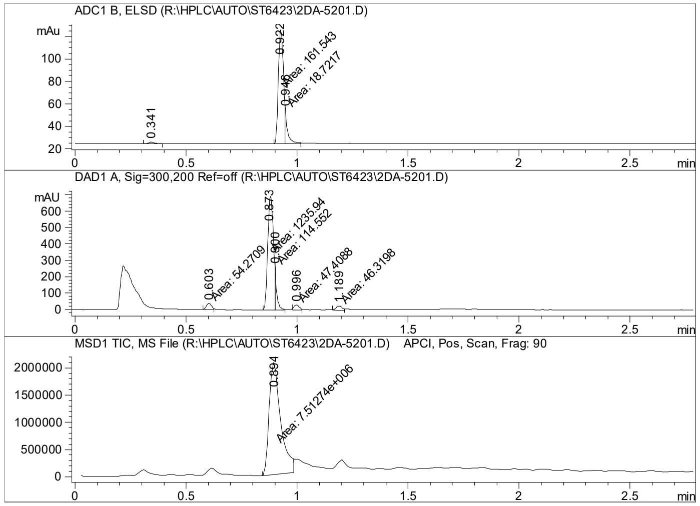


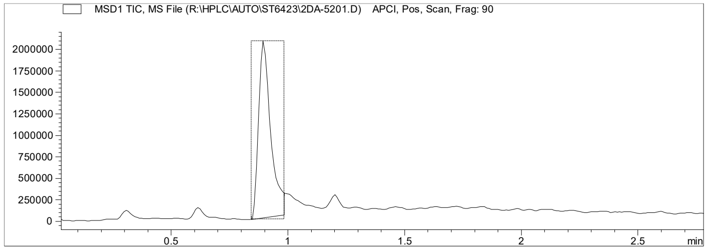


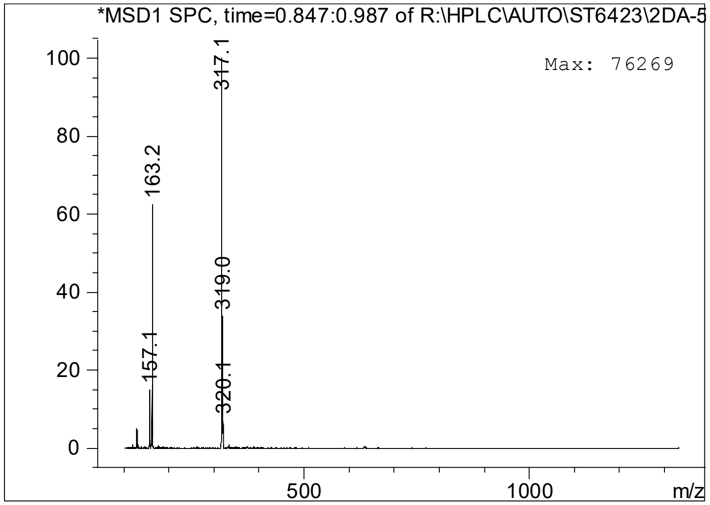

**LC-75**

**1-(4-bromophenoxy)-3-(1H-imidazol-1-yl)propan-2-ol**

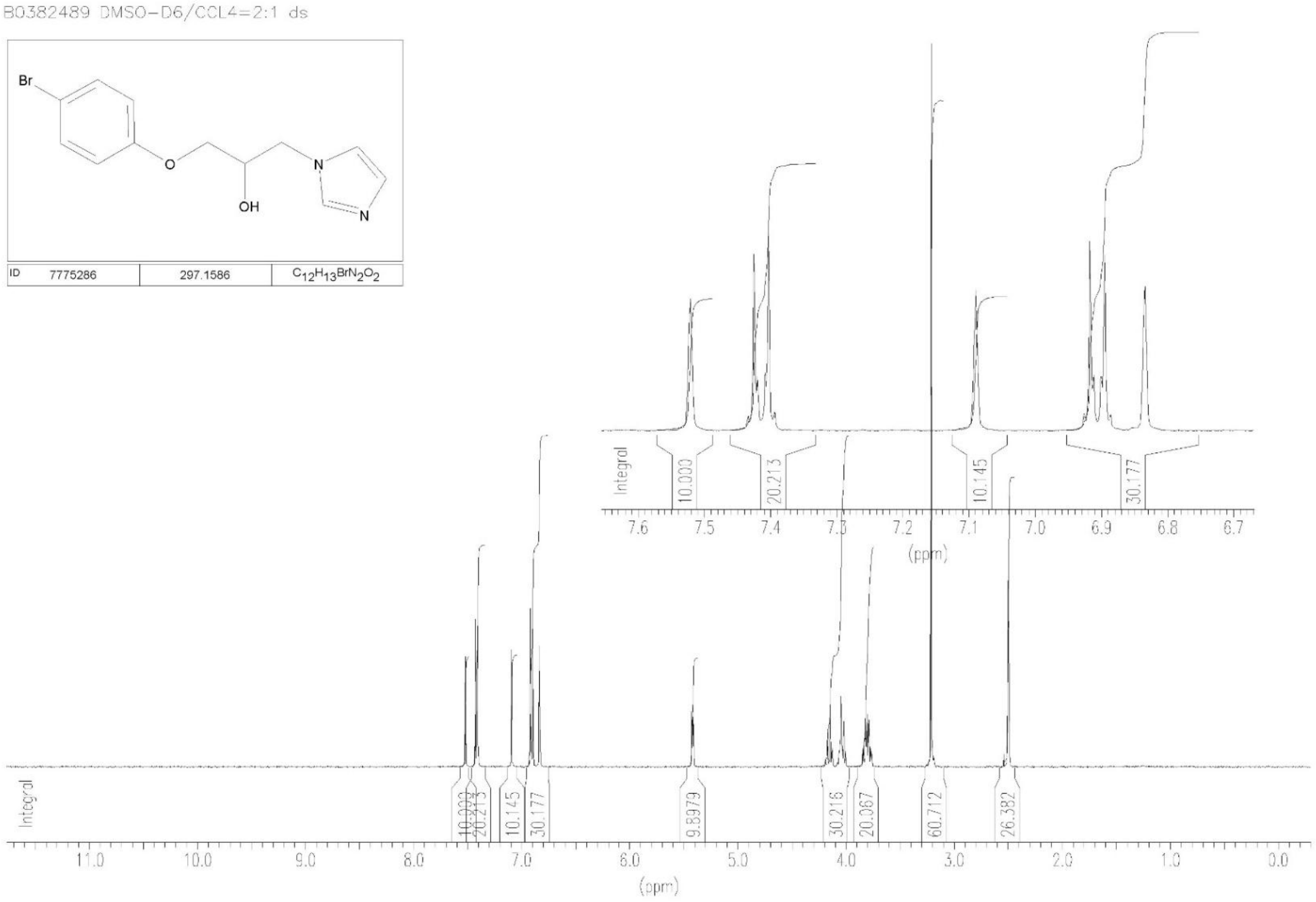
